# Organization of the *Drosophila* larval visual circuit

**DOI:** 10.1101/133686

**Authors:** Ivan Larderet, Pauline M. J. Fritsch, Nanaë Gendre, Larisa Neagu-Maier, Rick D. Fetter, Casey Schneider-Mizell, James W. Truman, Marta Zlatic, Albert Cardona, Simon G. Sprecher

## Abstract

Visual systems transduce, process and transmit light-dependent environmental cues. Computation of visual features depends on the types of photoreceptor neurons (PR) present, the organization of the eye and the wiring of the underlying neural circuit. Here, we describe the circuit architecture of the visual system of *Drosophila* larvae by mapping the synaptic wiring diagram and neurotransmitters. By contacting different targets, the two larval PR-subtypes create parallel circuits potentially underlying the computation of absolute light intensity and temporal light changes already within this first visual processing center. Locally processed visual information then signals via dedicated projection interneurons to higher brain areas including the lateral horn and mushroom body. The stratified structure of the LON suggests common organizational principles with the adult fly and vertebrate visual systems. The complete synaptic wiring diagram of the LON paves the way to understanding how circuits with reduced numerical complexity control wide ranges of behaviors.

## Introduction

Light-dependent cues from the surrounding world are perceived by specialized photoreceptor neurons (PRs) in the eye. Insect compound eyes are elaborate systems capable of mediating flight in a rapidly changing 3D environment. In contrast, larval stages present much simpler visual organs, which combined with their tractability, make them great models to link neural circuit processing and behavior (Kane et al., 2013; Randel et al., 2014, 2015; Gepner et al., 2015). Larvae of the fruit fly *Drosophila melanogaster* employ their visual system for a range of diverse behaviors including navigation, entrainment of circadian rhythms, formation of associative memories and may respond to the presence of other larvae (Kane et al., 2013; Humberg and Sprecher, 2017; Slepian et al., 20165; Justice et al., 2013; Yamanaka et al., 2013; von Essen et al., 2011; Gong, 2009; Mazzoni et al., 2005; Gerber et al., 2004; Sawin-McCormack et al., 1995). The simple eyes of the larva (also termed Bolwig Organ, BO) consist of only about 12 PRs each and yet drive a wide range of behaviors, raising questions on the organizational logic of the underlying visual circuit. Spectral sensitivity of PRs is defined by the *Rhodopsin* gene they express. Larval eyes contain two PR-types, either expressing the blue-tuned *Rhodopsin5* (Rh5) or the green-tuned *Rhodopsin6* (*Rh6*) (Malpel et al., 2002; Hassan et al., 2005; Rodriguez Moncalvo and Campos, 2005; Sprecher et al., 2007). Interestingly, for rapid navigation away from light exposure only the Rh5-PRs seem essential, whereas to entrain the molecular clock either PR subtype suffices (Keene et al., 2011). In the past, several neurons of the larval visual neural circuit have been identified but the logic of circuit wiring as well as the precise numbers of neurons involved in its first order visual processing center remain unknown.

Larval PRs project their axons in a joint nerve (Bolwig nerve) terminating in a small neuropil domain termed the larval optic neuropil (LON). Previous studies identified eleven neurons innervating the LON in each brain hemisphere. This includes four lateral neurons (LaNs) expressing the pigment dispersing factor (Pdf) neuropeptide (Pdf-LaNs) and a fifth non-expressing-Pdf LaN (5^th^-LaN), all being part of the clock circuit (Kaneko et al., 1997), as well as a serotonergic neuron and three optic lobe pioneer cells (OLPs) (Helfrich-Forster, 1997; Rodriguez Moncalvo and Campos, 2005, 2009; Tix et al., 1989). Recently, two unpaired median octopaminergic/tyraminergic neurons were described to extend neurites into these neuropils (Selcho et al., 2014). It remained unknown, however, whether these previously identified neurons constitute the entire neuronal components of the LON and how visual neuronal components connect to each other to form a functional network. In a recent study we started to investigate the anatomy of the LON using serial-section transmission electron microscopy (ssTEM) showing that PRs axons form large globular boutons with polyadic synapses and that the OLPs were parts of their direct targets (Sprecher et al., 2011).

Here, we mapped the synaptic wiring diagram of the LON by reconstructing all its innervating neurons from a new ssTEM volume of a whole first instar larval central nervous system (Ohyama et al., 2015). We characterized and quantified the connectivity of all previously described LON-associated neurons and identified new components in this circuit. We found per hemisphere eleven second order interneurons, two third-order interneurons and one serotonergic neuron innervating the LON, plus two unpaired octopaminergic/tyraminergic neurons contacting both hemispheres. We highlighted the separation of light signal flow at the first synapse level as the two PR subtypes connect onto distinct subsets of interneurons. Rh5-PRs connect to visual projection interneurons (VPNs) that transfer light information to distinct higher brain regions. Rh6-PRs synapse mainly on two visual local interneurons (VLNs) that appear to have key roles in light information processing by modulating the activities of Rh5-PRs targets. Network analysis suggests that VPNs may encode both absolute light intensity information (received from Rh5-PRs) and information about the changes of light intensity (received from the Rh6-PRs-VLNs pathway). These two VLNs are also the main targets of the aminergic neurons suggesting the modulation of the sensibility to light intensity variations by these central brain feedbacks. Comparison with the visual circuit of the adult fruit fly highlights common principles in organization of visual information processing, for example the stratification of PRs inputs as well as the existence of distinct photoreceptor pathways that are involved in detecting temporal light cues. Furthermore, the comparison with the olfactory wiring diagram (Berck et al., 2016) highlights common strategies for early sensory information processing, relay to higher order areas such as the mushroom bodies for associative memory, and control from the central brain. By mapping the connectivity of visual circuits and analyzing its architecture we have the opportunity to study the circut structure-function relationship and advance our understanding of how neural circuits govern behavior.

## Results

### Neurons of the larval visual circuit

Axonal projections of larval photoreceptor neurons (PRs) enter the brain lobes ventro-laterally via the Bolwig nerve and terminate in a small neuropil domain, termed larval optic neuropil (LON; Figure 1A,B, Figure 1-figure supplement 1). Visual interneurons innervate the LON from the central brain through the central optic tract (Sprecher et al., 2011). We reconstructed the axon terminals of all PRs and all their synaptic partners, as well as additional LON-innervating neurons that do not form synapses with PRs, from a serial-section transmission electron microscopy (ssTEM) volume spanning the complete central nervous system of a *Drosophila* first instar larva (Ohyama et al., 2015). In this way, we identified the complete repertoire of LON neurons and mapped the wiring diagram of the left and right LON.

**Figure 1:**
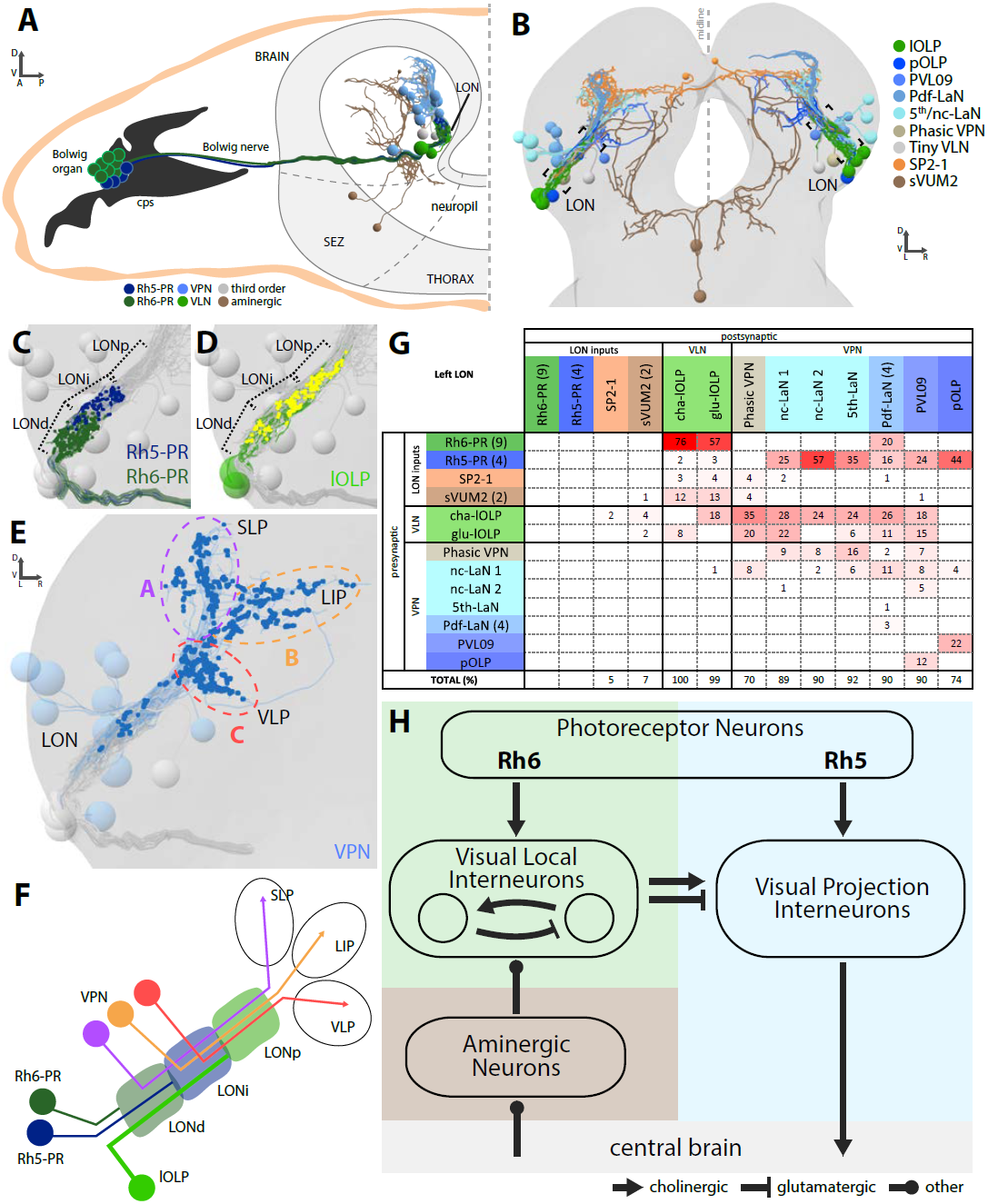
Overview of the larval optic neuropil. **A:** Schematic of the larval visual system with EM-reconstructed skeletons of all larval optic neuropil (LON) neurons. The Rh5-PRs (dark blue) and Rh6-PRs (dark green) cell bodies form the Bolwig organ sitting in the cephalopharyngeal skeleton (cps). They extend their axons to the brain via the Bolwig nerve. In the brain, neurons cell bodies are in the outer layer (gray) and project neurits into the neuropil. We can distinguish four main classes of neurons: visual projection interneurons (VPN, blue), visual local interneurons (VLN, green), third-order interneurons (gray) and aminergic modulatory neurons (brown). Octopaminergic/tyraminergic modulatory neurons cell bodies sit in the subesophageal zone (SEZ). **B:** 3D reconstruction of all LON-associated neurons from the ssTEM dataset in both hemispheres (except Bolwig nerves): VLN in green: local optic lobe pioneer neurons (lOLPs); VPN in shades of blue: the projection OLP (pOLP), a novel neuron which is located in the posterior ventral lateral cortex (PVL09), the Pdf-lateral neurons (Pdf-LaNs), the 5th-LaN and the non-clock-LaNs (nc-LaNs); third-order neurons: Phasic VPN in light brown and Tiny VLN in gray; aminergic modulatory neurons: serotonergic neuron (SP2-1, orange) and SEZ-ventral-unpaired-medial-2 octopaminergic/tyraminergic neurons (sVUM2, brown). Posterior view. **C-E:** 3D representations of the presynaptic sites of LON neurons in the left lobe, posterior view. C: Rh6-PRs presynaptic terminals (dark green) define a distal LON layer (LONd) while Rh5-PRs presynaptic connections (dark blue) define an intermediate LON layer (LONi). A third layer of the LON, more proximal (LONp) is devoid of PR terminals. Other LON neurons in gray. **D:** All LON layers, including the LONp, contain presynaptic sites from the lOLPs (skeletons in green, synapses in yellow). VPNs, Tiny VLN and Bolwig nerve in gray. **E:** VPNs (blue) make synaptic connections in three main regions outside the LON. VPNs projections define three domains: dorsal domain (A, violet) defined by projections in the superior lateral protocerebrum (SLP), lateral domain (B, orange) in the lateral inferior protocerebrum (LIP), ventral domain (C, red) in the ventral lateral protocerebrum (VLP). VLNs and Bolwig nerve in gray. **F:** Schematic of the LON three layers: LONd innervated by Rh6-PRs (dark green), LONi innervated by Rh5-PRs (dark blue) and LONp innervated by lOLPs (green); and of the three domains outside the LON were different VPNs subtypes project to (violet, orange and red empty circles). lOLPs also make presynaptic connections in the LONd and LONi (thick line). **G:** Connectivity table of the left LON with the percentage of postsynaptic sites of a neuron in a column from a neuron in a row. Neurons of same type are grouped, in brackets number of neurons in the group. Same colors as in B. Only connections with at least two synapses found in both hemispheres were used. **H:** Simplified diagram of the larval visual system. PRs neurons inputs are cholinergic and define two pathways. Rh5-PRs target VPNs (blue area) while Rh6-PRs target the two main larval VLNs (green area). Between these two VLNs, one is cholinergic while the other one is glutamatergic and they both inputs onto VPNs. These VLNs also integrate aminergic modulatory inputs (brown area) that potentially bring information from the central brain. VPNs project to higher order regions of the brain (gray area).

We define five neuron types (Figure 1A,B): first, sensory neurons (photoreceptor neurons) that innervate the LON; second, visual local interneurons (VLNs) that do not extend neurites beyond the LON; third, visual projection interneurons (VPNs) that relay signals from the LON to distinct higher brain areas; fourth, third-order interneurons in the LON that do not receive direct input from the PRs; and fifth, modulatory aminergic feedback neurons projecting from the central brain.

Two interneurons belonging to the previously described optic lobe pioneer cells (OLPs, Tix et al., 1989) are VLNs, which we therefore named local-OLPs (lOLPs, Figure 2). Their arbors are fully contained within the LON and they present a distinct axon and dendrite (Figure 2B,C), comparable to glutamatergic inhibitory neurons of the larval antennal lobe (Berck et al., 2016). We found that one lOLP is cholinergic (cha-lOLP) while the other is glutamatergic (glu-lOLP), in agreement with previous studies (Figure 2-figure supplement 1; Yasuyama et al., 1995; Daniels et al., 2008). Whereas acetylcholine is known to act as an excitatory neurotransmitter in *Drosophila* (Baines and Bate, 1998; Burrows 1996), glutamate has been found to decrease Pdf-LaNs calcium level (Hamasaka et al. 2007) and to mediate inhibition in the antennal lobe (Liu and Wilson, 2012; Berck et al. 2016), suggesting that the glulOLP VLN is putatively inhibitory, a role consistent with its position in the visual circuits (see discussion).

**Figure 2:**
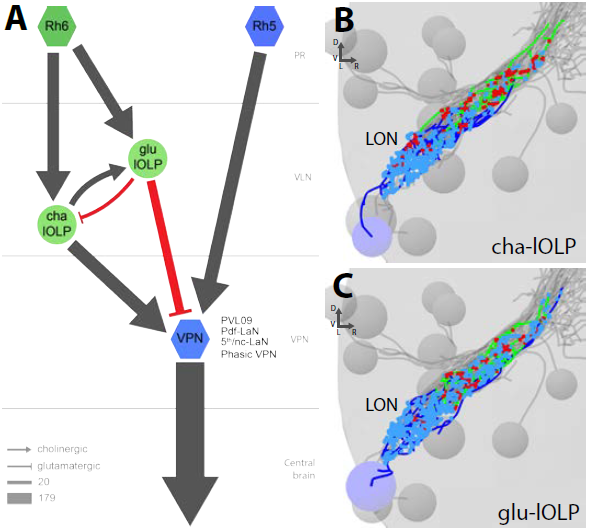
The two main VLNs of the larval visual system. **A:** Wiring diagram of both lOLPs (light green circles). The two lOLPs receives from Rh6-PRs (dark green) and are reciprocally connected, suggesting feedback inhibition from the glu-lOLP onto the cha-lOLP. They also share the same type of targets: VPNs (blue) including PVL09, all LaNs and the Phasic VPN, that are direct targets of Rh5-PRs (dark blue) (except the Phasic VPN) and are outputs of the LON towards the central brain. Left hemisphere, hexagons represent group of cells, circles represent single cell, arrow thickness weighted by the square root of the number of synapses, arrow thickness scale shows minimum and median. **B-C:** 3D reconstructions of both lOLPs (cha-lOLP (**B**) and glu-lOLP (**C**)) with dense arborizations within the LON. Posterior view, dendrites in blue, axons in green, presynaptic sites in red, postsynaptic sites in cyan, other LON neurons in gray.

VPNs (Figure 3) include the previously identified pigment dispersing factor (Pdf)-expressing lateral neurons (Pdf-LaNs) and the Pdf-negative 5^th^-LaN of the circadian clock circuit (Kaneko et al., 1997), as well as one neuron belonging to the OLPs (Tix et al., 1989) which we named accordingly projection-OLP (pOLP). In addition, in the VPN group we newly identified two non-clock lateral neurons (nc-LaNs) originating from the same neuroblast lineage as the 5^th^-LaN (Figure 3I; Figure 3-figure supplement 1), and one neuron defined by its “postero-ventro-lateral” cell body position, termed PVL09. All these VPNs, except the four peptidergic Pdf-LaNs, are cholinergic (Figure 3-figure supplement 1).

**Figure 3:**
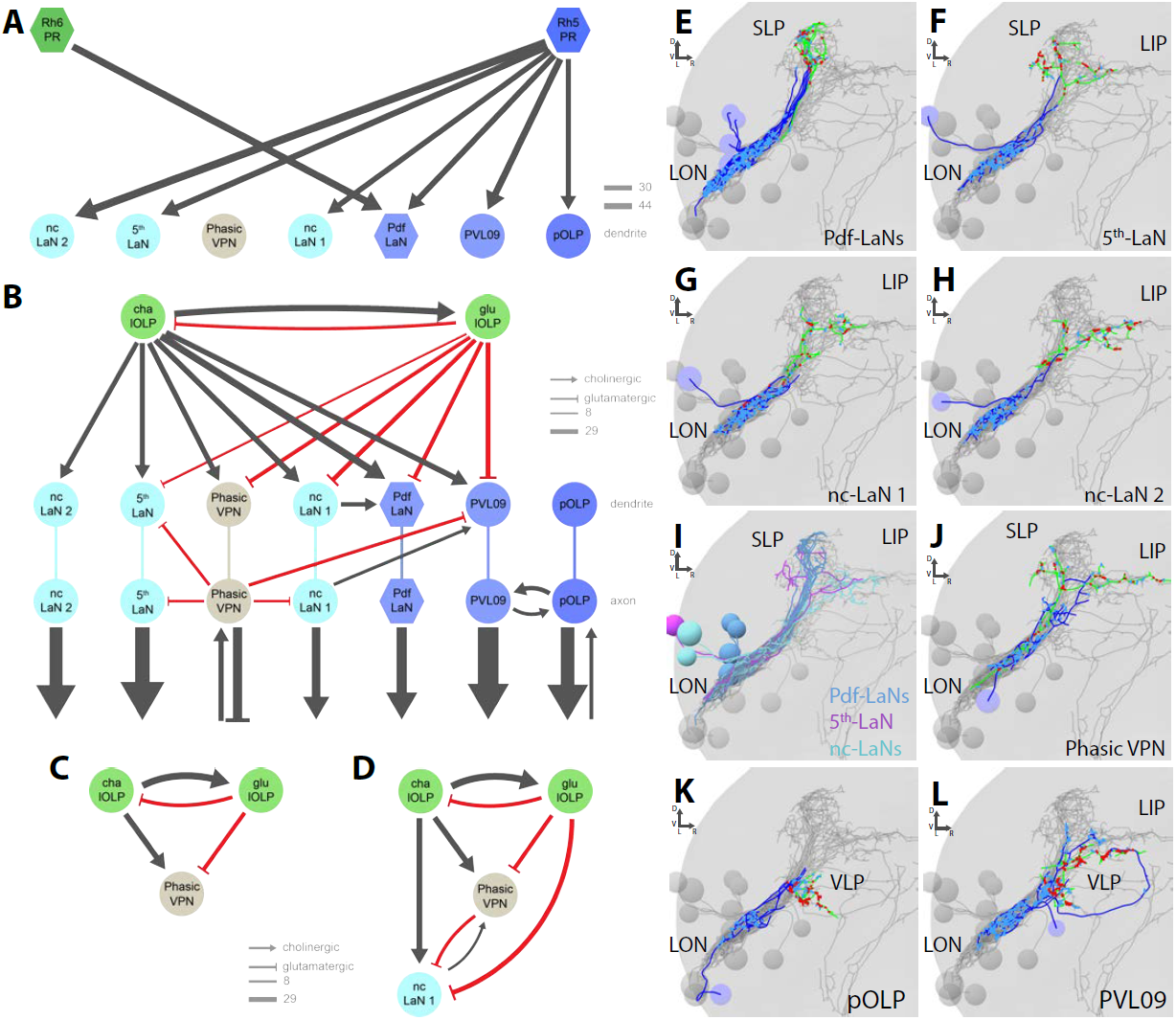
VPNs of the visual system. **A-D:** Wiring diagrams of VPNs (shades of blue for Rh5-PRs targets plus the Phasic VPN in light brown). Left hemisphere, hexagons represent group of cells, circles represent single cell, arrow thickness weighted by the square root of the number of synapses, arrow thickness scale shows minimum and median. **A:** The four Pdf-LaNs are the only VPNs that receive from both Rh6-PRs (dark green) and Rh5-PRs (dark blue). The Phasic VPN is a third-order neuron that does not receive any inputs from PRs. All other VPNs receive visual inputs uniquely from Rh5-PRs. All inputs from PRs onto VPNs are situated on the target dendrites. **B:** VPNs except pOLP are targets of the two lOLPs (light green) and these connections are situated on the VPNs dendrites. Additionally, PVL09 receives inputs from both the Phasic VPN and nc-LaN 1 while the Pdf-LaNs receive only from nc-LaN 1, and the 5th-LaN receive only from the Phasic VPN. PVL09 and pOLP are reciprocally connected at their axon level. All VPNs transfer light information to neurons deeper in the brain. The Phasic VPN and pOLP additionally receive on their axons some inputs from other neuronal circuits. **C:** Circuit motif of the Phasic VPN receiving from both lOLPs. **D:** Circuit motif of the nc-LaN 1 that is under excitation from cha-lOLP, tonic inhibition from glu-lOLP and phasic inhibition from the Phasic VPN. Moreover nc-LaN 1 connects back to the Phasic VPN allowing a possible OFF response motif. Similar motifs can be described for other VPNs (Figure 3-figure supplement 3). **E-L:** 3D reconstructions from ssTEM dataset, posterior view, dendrites in blue, axons in green, presynaptic sites in red, postsynaptic sites in cyan, other LON neurons in gray. VLP: ventral lateral protocerebrum. SLP: superior lateral protocerebrum. LIP: lateral inferior protocerebrum. **E:** The four Pdf-LaNs project to the SLP. **F:** The 5th-LaN projects both to the SLP and the LIP region, whereas nc-LaN 1 and 2 (**G** and **H**) mainly project to the LIP. **I:** Anatomy of all LaNs together. **J:** The Phasic VPN cell body is situated anteriorly to the LON and it has an axon coming back in the LON in top of its projections within both SLP and LIP regions. **K:** pOLP cell body is situated with the lOLP and projects to the VLP. **L:** PVL09 cell body is situated postero-ventro-laterally to the LON and has an axon with a characteristic loop shape, extending first towards the ventro-medial protocerebrum, then towards the LIP before curving down back to the VLP, where it forms most of its synaptic output.

We also identified two third-order interneurons that make connections within the LON but do not receive direct inputs from PRs. The first one, that we named Phasic VPN according to its suggested function (see discussion), is defined by prominent axonal projections beyond the LON and significant pre-synaptic termini within the LON (Figure 3J). We found that the Phasic VPN is glutamatergic and therefore putatively inhibitory as well (Figure 3-figure supplement 1). The second third-order interneuron is a VLN that we named Tiny VLN because of its small size in the current dataset (Figure 1-figure supplement 4; no additional information could be collected as no known GAL4 line labels this cell).

Finally, LON circuits are modulated from the central brain by a bilateral pair of serotonergic neurons and two ventral-unpaired-medial neurons of the subesophageal zone that are octopaminergic/tyraminergic and that project bilaterally to both LON (Figure 4; Huser et al., 2012; Rodriguez Moncalvo and Campos, 2009; Selcho et al., 2014). These neurons match in number and neuromodulator type with the left-right pair of serotonergic neurons and the two bilateral octopaminergic neurons of the larval antennal lobe (Berck et al. 2016), providing support for an ancestral common organization of the visual and the olfactory sensory neuropils (Strausfeld 1989; Strausfeld et al. 2007).

**Figure 4:**
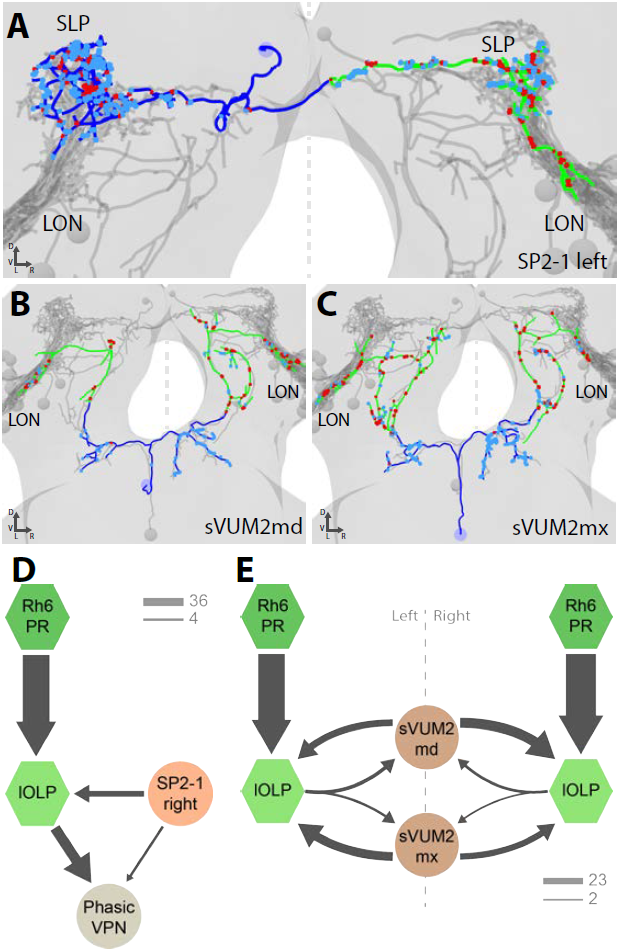
Aminergic modulatory inputs of the larval visual system. **A-C:** 3D reconstructions from ssTEM data, posterior view, dendrites in blue, axons in green, presynaptic sites in red, postsynaptic sites in cyan, other LON neurons in gray, dashed line represent brain midline. **A:** The SP2-1 neuron from the left hemisphere innervates the ipsilateral SLP and the contralateral LON. sVUM2md (**B**) and sVUM2mx (**C**) neurons are located along the midline in the SEZ with their neurit splitting and innervating both hemispheres in a symmetric fashion. Their bilaterally symmetrical branches receive synaptic input in the SEZ and extend their axon towards the protocerebrum prior to turning laterally and entering the LON. Branches within the protocerebrum and LON contain presynaptic and postsynaptic sites. **D:** Connectivity graph showing the SP2-1 neuron (orange) of the right hemisphere connecting with the lOLPs (light green) and the Phasic VPN (light brown) of the left hemisphere. Connections between the lOLPs and the Phasic VPN are also displayed as well as lOLPs inputs from Rh6-PRs (dark green). **E:** Connectivity graph of sVUM2mx and sVUM2md (brown) showing that their only partners are the lOLPs (light green) but in both hemispheres. D-E: Hexagons represent group of cells, circles represent single cell, arrow thickness weighted by the square root of the number of synapses, arrow thickness scales shows minimum and median.

With the exception of the unpaired octopaminergic/tyraminergic neurons, we identified in all cases pairs of bilaterally homologous VLNs and VPNs. In addition, in the right brain hemisphere we found an additional fourth OLP, which, together with the variable number of PRs, suggests that the circuit architecture can accommodate a variable number of neurons (Figure 5; see below).

**Figure 5:**
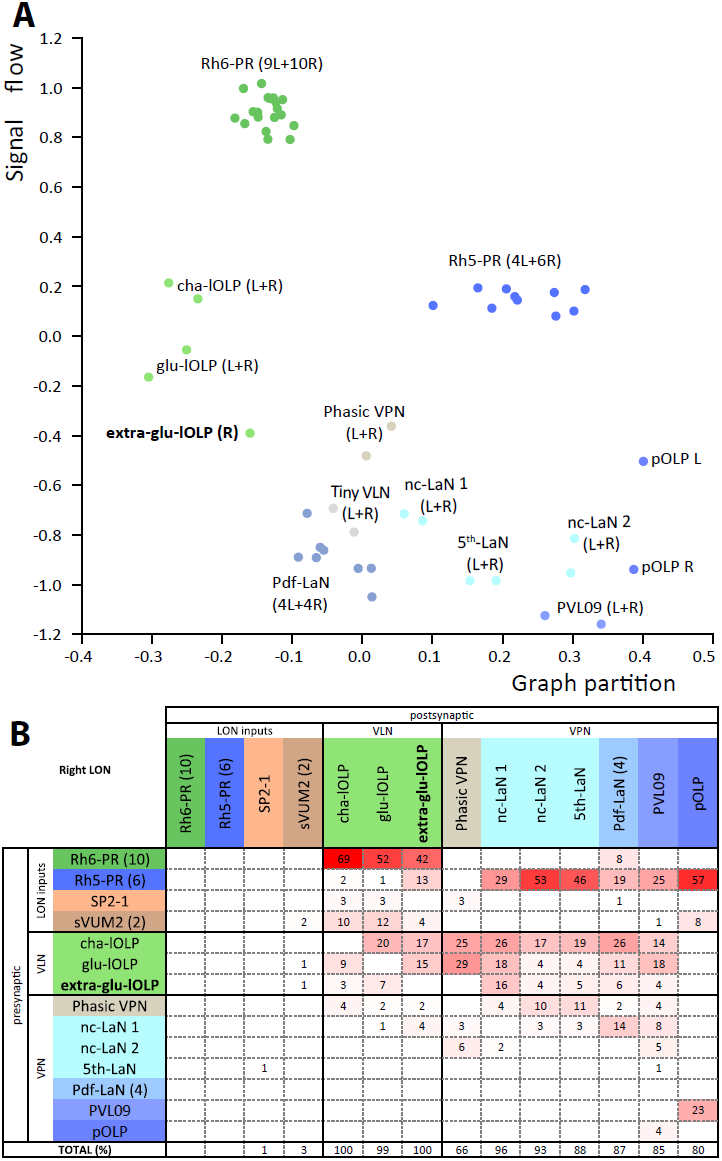
Larval optic neuropil architecture is maintained despite a variable number of neurons. Color code as in Figure 1. **A:** Graph partition of the visual neurons from the left (L) and right (R) hemispheres. We excluded the neuromodulatory neurons SP2-1, which are weakly connected, and the sVUM2md and sVUM2mx, which project bilaterally. Visual information flows from PRs at the top towards VPNs at the bottom (Varshney et al., 2011). Note how the extra-glu-lOLP of the right hemisphere (bold) positioned closely with the other lOLPs. **B:** Connectivity table of the right LON with the percentage of postsynaptic sites of a neuron in a column from a neuron in a row. Only connections with at least two synapses found in both hemispheres were used (except for the extra-glu-lOLP).

### The larval optic neuropil is organized in three layers

Visual circuits in the mammalian retina as well as in the optic ganglia of the adult fruit fly are organized in layers. These layers are characterized by dendritic arborizations or axonal termini of specific neuron types (Sanes and Zipursky, 2010). In *Drosophila* larvae, the LON can be subdivided into three distinct layers based on the innervation of the PR subtypes (Sprecher et al., 2011). Briefly, Rhodopsin6-PRs (Rh6-PRs) terminate in the distal, most outer layer of the LON (LONd), whereas Rhodopsin5-PRs (Rh5-PRs) terminate in the intermediate LON layer (LONi). The most proximal, inner layer of the LON (LONp) lacks direct PRs input (Figure 1C,D,F).

The layered arrangement of PRs axon terminals translates into specific connectivity with LON neurons. Most VPNs, whose dendrites do not reach the LONd, receive direct inputs from Rh5-PRs only, whereas the dendrites of Pdf-LaNs span both the LONd and LONi, integrating inputs from both Rh5-and Rh6-PRs (see below; Figure 3E-L). The absence of PRs axon terminals in the LONp deprives the third-order interneurons, Phasic VPN and Tiny VLN, whose dendrites are restricted to the LONp, from any direct PRs inputs (*Figures 1G, 3J*). Intriguingly, the Tiny VLN integrates inputs within the LONp and projects back to both LONi and LONd (Figure 1-figure supplement 4).

Beyond the LON, VPNs axons target three distinct protocerebral areas, namely the superior lateral protocerebrum, the lateral inferior protocerebrum and the ventro lateral protocerebrum (Figure 1E,F). Interestingly, these areas overlap in parts with the lateral horn, involved in innate behaviors, and the mushroom body calyx, involved in associative memory (see below). Within the LON, the dendrites of VPNs are mainly postsynaptic, whereas their axons, upon reaching higher brain areas, present both presynaptic and postsynaptic sites (Figure 3C-J). This suggests that VPNs outputs are modulated by input from non-visual neurons, similarly to how olfactory projection interneurons receive non-olfactory inputs (Berck et al., 2016). In particular pOLP and the Phasic VPN receive up to 30% of their inputs from non-LON neurons (Figure 1G).

### Parallel light input channels: each PR subtype targets distinct VPNs and VLNs

Previous studies suggested that only Rh5-PRs are critical for rapid light avoidance, while Rh6-PRs appeared non-essential (Keene et al., 2011; Kane et al., 2013). However, for entrainment of the molecular clock either PR-type by itself is sufficient (Keene et al., 2011). These findings lead to speculate that Rh5-PRs and Rh6-PRs connect to distinct types of visual interneurons. Supporting this notion, we found that Rh6-PRs synapse principally onto VLNs (79%) and much less onto VPNs (15%, all onto one single VPN type: the Pdf-LaNs; Figure 1G). Conversely, Rh5-PRs preferentially synapse onto VPNs (90%) and much less onto VLNs (6%). Pdf-expressing LaNs of the clock circuits are the only interneurons that receive direct inputs from both PR-subtypes (20% from Rh6-PRs and 15% from Rh5-PRs, Figure 1G), supporting behavioral evidence that either PR-subtype may entrain the larval clock (Keene et al., 2011).

Rh6-PRs target the two main VLNs of the LON: the cha-and glu-lOLP (*Figures 1G, 2A*). Furthermore, the lOLPs main inputs come from Rh6-PRs: up to 75% of cha-OLP input and 58% of glu-lOLP input. Importantly, these two VLNs synapse onto most VPNs including the four Pdf-LaNs and the 5^th^-LaN of the circadian clock, the nc-LaNs, PVL09 and the third-order interneuron Phasic VPN (Figure 3B). pOLP is the only VPN that does not receive inputs from its sister cells, the two lOLPs, and therefore create a direct output pathway of the Rh5-PRs light dependent information towards higher brain regions (Figure 3A, B). pOLP is strongly interconnected through axo-axonic connections with PVL09 suggesting that they may reciprocally cross-enhance their excitatory synaptic output (*Figures 1G, 3B;* Figure 1-figure supplement 3). Also, the 5^th^-LaN and the two nc-LaNs, present different fractions of inputs from the glu-lOLP, suggesting that they will encounter different levels of postsynaptic inhibition from this cell (see discussion, *Figures 1G, 3B*).

The third-order Phasic VPN, which does not receive direct inputs from PRs (Figure 3A), is downstream of the two lOLPs and is itself connecting onto several VPNs including the 5^th^-LaN, the two nc-LaNs and PVL09 creating another layer of possible computation (see discussion, Figure 3B, C, D).

In summary, most VPNs that directly integrate Rh5-PRs light dependent information may be modulated indirectly by the Rh6-PRs light dependent information via the two VLNs, cha-and glu-lOLPs (*Figures 1H, 2A*).

### VPNs target different brain areas

Distinct areas of the protocerebrum are innervated by the six unique VPNs of the larval visual circuit (the pOLP, the 5^th^-LaN, the two nc-LaNs, the third-order interneuron Phasic VPN, and PVL09) and by the four Pdf-LaNs of the circadian clock.

pOLP targets the lower lateral horn, an area also innervated by the multiglomerular olfactory projection interneuron (mPN) Seahorse (Berck et al., 2016; Figure 3K, Figure 3-figure supplement 2) with whom it shares numerous postsynaptic partners (data not shown). Since mPN Seahorse integrates inputs from the aversive OR82a-expressing olfactory receptor neuron (Kreher et al., 2008), downstream neurons of pOLP and mPN Seahorse are likely contributing to aversive behavior.

Three sister VPNs (the 5^th^-LaN and both nc-LaN 1 and 2) present similar axon trajectories, dropping synapses in the lateral horn until reaching the accessory calyx of the mushroom body (Figure 3F-H, Figure 3-figure supplement 2), where they synapse onto Kenyon cells (Eichler et al., 2017). On top of its local connections and potential function in phasic inhibition (see discussion), the Phasic VPN also has projections beyond the LON in a similar pattern as both nc-LaNs (Figure 3J, Figure 3-figure supplement 2).

PVL09 is unique among the VPNs in presenting a bifurcated axon with one branch following the other VPNs into the lateral horn and the other branch taking a long looping path below the mushroom body before coming back to the same region as pOLP (Figure 3L, Figure 3-figure supplement 2). Like most VPNs, PVL09 is under the control of the two lOLPs and the Phasic VPN, but additionally receives inputs from both nc-LaNs on its dendrites and axon (Figure 3-figure supplement 3; Figure 1-figure supplement 3), suggesting that it integrates broadly all light information.

In summary, all six unique VPNs form synapses in the lateral horn, and of the four VPNs (all but pOLP and Phasic VPN) that potentially encode a mixture of absolute light intensity and changes in light intensity (see discussion), three (the 5^th^-LaN, nc-LaN 1 and nc-LaN 2) synapse onto both the lateral horn and the mushroom body Kenyon cells (Eichler et al., 2017). VPN connections onto Kenyon cells may underlie the larval ability to form associative memories with light as a conditioned stimulus, whereas light as an unconditioned stimulus could be encoded via their connections onto the lateral horn (von Essen et al., 2011).

Finally, the Pdf-LaNs, necessary for circadian rhythm, project to a region dorsal and more medial to the lateral horn, similarly as in adult *Drosophila* (Yasuyama and Meinertzhagen, 2010), where they make few small dyadic synapses from boutons rich in dense-core vesicles (Figure 3E, Figure 3-figures supplement 1 and 2). Pdf-LaNs boutons also contain clear vesicles suggesting that they might co-express a neurotransmitter, which in adult flies has been suggested to be glycine (Figure 4-figures supplement 1; Frenkel et al., 2017).

### Central brain feedback via octopaminergic/tyraminergic and serotonergic neurons

Similarly to other sensory modalities (Roy et al., 2007; Dacks et al., 2009; Huser et al., 2012; Selcho et al., 2014; Majeed et al., 2016; Berck et al., 2016), a set of aminergic neurons provide feedback from the central brain into the LON, thus creating an entry point to modulate visual information processing. Both types of modulatory neurons have previously been identified in the LON (Rodriguez Moncalvo and Campos, 2005; Huser et al., 2012; Selcho et al., 2014).

A pair of serotonergic neurons belonging to the SP2 cluster (named SP2-1) connects to the contralateral LON, while receiving presynaptic input predominantly in the ipsilateral protocerebrum (Figure 4A, Figure 4-figure supplement 1). The two other aminergic input neurons are the octopaminergic/tyraminergic subesophageal zone-ventral-unpaired-medial 2 neurons of the maxillary and mandibular clusters (sVUM2mx and sVUM2md, Figure 4B,C, Figure 4-figure supplement 1). Each sVUM2 innervates both brain hemispheres in a symmetric fashion.

Each SP2-1 neuron connects to the main VLNs (cha-and glu-lOLP) as well as to the third-order Phasic VPN of the contralateral side (Figure 4D). The octopaminergic/tyraminergic sVUM2 neurons uniquely synapse onto the two lOLPs but in both hemispheres simultaneously (Figure 4E). In contrast to the SP2-1, the lOLPs form feedback synapses onto the axonal termini of both sVUM2 neurons. This feedback motif may allow local tuning of the octopaminergic/tyraminergic modulatory input, whereas the serotonergic input is not altered within the LON.

In summary, SP2-1 and sVUM2 mediate feedback from other brain areas to potentially modulate the sensitivity to light fluctuations (see discussion). ON and OFF responses may be further affected by inputs from SP2-1 onto the Phasic VPN (see discussion). These possible modulations arise from monosynaptic connections between the aminergic cells with the lOLPs and the Phasic VPN, while additional effects might be elicited by volume release of serotonin and octopamine (Dacks et al., 2009, Linster and Smith, 1997, Selcho et al., 2012). Further reconstruction is needed to identify the presynaptic inputs of these aminergic modulatory neurons.

### Bilaterally symmetric LON circuits with asymmetric numbers of neurons

Similarly to the non-stereotypic number of ommatidia in the compound eye of the adult fly (Ready et al., 1976), the precise number of PRs in each larval eye also varies (Sprecher et al., 2007). In the current specimen, we identified thirteen PRs in the left hemisphere and sixteen in the right hemisphere. We found four Rh5-PRs and nine Rh6-PRs in the left hemisphere, and six Rh5-PRs and ten Rh6-PRs in the right hemisphere. Despite this difference in PRs number, homologous LON interneurons in each hemisphere receive a similar fraction of inputs from PRs (Figure 5-figure supplement 1), similarly to olfactory projection interneurons that receive an equivalent fraction of inputs from olfactory receptor neuron despite differences in the numbers of these sensory neurons (Tobin et al., 2017). This further supports the idea that projection interneurons may regulate the amount of inputs from sensory neurons that they receive relative to the total inputs on their dendrites.

Interestingly, we found a third local-OLP in the right brain hemisphere. Similarly to a non-stereotypic PRs number, variability in OLPs number has been observed before (Tix et al., 1989). When using a GAL4 driver labeling glutamatergic neurons (OK371; Mahr and Aberle, 2005) we found an extra-glutamatergic-OLP (Figure 5-figure supplement 1) at a similar frequency as the presence of the fourth OLP cell had been reported (in about 5% of brains) and displaying an asymmetry between hemispheres. It is unusual to have variability and asymmetries in *Drosophila* neural circuits (Ohyama et al., 2015; Berck et al., 2016; Schlegel et al., 2016; Jovanic et al., 2016; Schneider-Mizell et al., 2016) but it has been observed before (Takemura et al., 2015; Tobin et al., 2017; Eichler et al., 2017). The presence of an extra-glu-lOLP and variable numbers of PR raises the question of the overall stereotypy of circuit architecture when comparing the left and right LON.

We analyzed the structure of the left and right LON circuits with spectral graph analysis of the connectivity matrices by plotting the graph partition metric as a function of the signal flow metric (Varshney et al., 2011). Despite the left and right LON not sharing any interneurons and having a different number of PR and lOLP cells, we observed that neurons of the same type cluster closely together (Figure 5A), indicating that the circuit structure in which each identified cell is embedded is very similar in the both hemispheres. The position of the extra-lOLP in this spectral graph analysis plot and its choice of pre-and postsynaptic partners (Figure 5B), in particular the many inputs from Rh6-PRs and cha-lOLP, suggest that the extra-lOLP may act as an extra-glu-lOLP, in agreement with its inclusion in the OK371-GAL4 expression pattern. Its reciprocal connections with the Tiny VLN are also in favor of this hypothesis (Figure 1-figure supplement 4). Note though that this extra-glu-lOLP also receives some inputs from Rh5-PRs and lacks inputs from the serotonergic neuron unlike other lOLPs, which indicates that it might still act differently than a glu-lOLP. Why some larvae present this additional cell remains to be determined. Importantly, connections among other LON neurons do not seem affected by the presence of an extra-glu-lOLP (Figure 5B; Figure 5-figure supplement 1). In conclusion, while the number of PRs and VLNs can vary in the larval visual system, the overall circuit architecture is maintained, and in particular the output channels (VPNs) are identical.

## Discussion

A shared characteristic of many visual systems is the retinotopic organization allowing visual processing in a spatially segregated fashion by the transformation of the surrounding environment into a 2D virtual map (for review, Sanes and Zipursky, 2010). Within the compound eye of the adult fruit fly, this is achieved by the prominent organization of PRs in ommatidia in the retina that is maintained through underlying cartridges in the lamina and columns in the medulla. The *Drosophila* larval eye lacks ommatidia or a similar spatial organization of PRs. Nevertheless larvae can navigate directional light gradients and form light associative memories using their simple eyes (Kane et al., 2013; von Essen et al., 2011; Humberg and Sprecher, 2017).

In this study, we described the synapse-level connectome of the larval first visual center by reconstructing neurons recursively from the optic nerves to third-order neurons following all chemical synapses in a nanometer-resolution EM volume of the whole central nervous system. We found that the two PR subtypes synapse onto distinct target interneurons, showing a clear separation of visual information flow from either Rh5-PRs or Rh6-PRs. Rh5-PRs predominantly synapse onto VPNs, which transfer light information to distinct regions including the lateral horn and the mushroom body calyx, whereas Rh6-PRs strongly synapse onto two VLNs (cha-and glu-lOLP). Moreover, the flow of information is convergent as these two main VLNs in turn synapse onto most VPNs. These two main VLNs also receive input from both the serotonergic as well as the octopaminergic/tyraminergic systems, suggesting that their activity may be modulated by input from central brain circuitry. Thus, a key feature of the larval visual circuit is that the Rh6-PRs-pathway feeds into the Rh5-PRs-pathway, suggesting a tuning function for Rh6-PRs and the two VLNs that also integrate external modulatory inputs (Figure 1H; Figure 1-figure supplement 5). This is not excluding the possibility of electrical connections mediated by gap junctions that are not visible in the ssTEM volume.

### The Rh6-PRs-VLNs pathway may compute variations in light intensity

In other sensory systems such as the olfactory system of the adult (Rybak et al., 2016) and larval *Drosophila* (Berck et al., 2016) and in the chordotonal mechanosensory systems of the locust (Wolff and Burrows, 1995) and *Drosophila* larva (Ohyama et al., 2015, Jovanic et al., 2016) axonal terminals of sensory neurons receive abundant inhibitory inputs from central neurons. Such presynaptic inhibition of sensory terminals may be employed to mediate lateral inhibition (Wilson and Laurent, 2005; Olsen and Wilson, 2008), to implement divisive normalization (Olsen et al. 2010) or to encode a predicted future stimulation (Wolf and Burrows, 1995). However, in the larval visual circuit, we do not observe significant synaptic connections onto the PRs (*Figures 1G, 5B*), suggesting that the light information encoded by PRs is passed on with no alteration at the level of the first synapse.

However, as mentioned above, the Rh6-PRs inputs are relayed to the VPNs only via the two VLNs, the cha-and glu-lOLPs. Considering cha-lOLP as excitatory and glu-lOLP as putatively inhibitory, Rh6-PRs signals are positively transferred to VPNs only via the cha-lOLP (Figure 2A). In addition, cha-lOLP receives strong inputs from the putatively inhibitory glu-lOLP (*Figures 1G, 2A*), which is also almost exclusively driven by Rh6-PRs inputs. Therefore, the strong putative inhibition of glu-lOLP onto cha-lOLP may mediate a form of indirect presynaptic inhibition of the Rh6-PRs inputs. This motif made by the cholinergic Rh6-PRs driving both the cha-lOLP and the glu-lOLP, and with the glu-lOLP putatively inhibiting the cha-lOLP, creates an incoherent feedforward loop (Alon, 2007). Therefore this motif could make the cha-lOLP responsive to the derivative of Rh6-PRs activity, i.e. to the variations in light intensity. As cha-lOLP VLN is likely to be a positive relay of Rh6-PRs inputs onto VPNs, VPNs could therefore respond to increments in light intensity.

Moreover, cha-lOLP inputs onto the glu-lOLP and both cha-and glu-lOLPs further connect onto the VPNs. Therefore, this creates a second incoherent feedforward motif making VPNs potentially responsive to the derivative of cha-lOLPs inputs. As cha-lOLPs inputs may already represent the first derivative of Rh6-PRs inputs, this second incoherent feedforward motif may therefore make VPNs responsive to the acceleration of light intensity raises.

In summary, glu-lOLP may provide both presynaptic inhibition of Rh6-PRs inputs by inhibiting the relay neuron cha-lOLP, and postsynaptic inhibition by directly inhibiting most VPNs. Consequently, VPNs could respond to the absolute light intensity (from Rh5-PRs) and to the variations of light intensity (from the Rh6-PRs-VLNs pathway). Interestingly, olfactory projection interneurons in the adult *Drosophila* also respond to a mixture of absolute odorant concentration and of the acceleration of odorant concentration (Kim et al., 2015). As for other sensory system (Klein et al., 2015; Schulze et al., 2015), a measure of changes in light intensity over time would enable light gradient navigation (Kane et al., 2013; Humberg and Sprecher, 2017).

### Phasic inhibition could sharpen ON and OFF responses

In the wiring diagram, we found a glutamatergic third-order interneuron, named Phasic VPN, mainly driven by cha-lOLP and therefore potentially only active upon an increase in light intensity via Rh6-PRs (Figure 3C). This is unlike the other glutamatergic interneuron, glu-lOLP, which receives direct inputs from PRs and therefore can be tonically active in the presence of light. The Phasic VPN specifically synapses onto multiple VPNs (the 5^th^-LaN, the two nc-LaNs and PVL09) and therefore potentially inhibits these cells that also receive strong input from glu-lOLP (Figure 3B, D, Figure 3-figure supplement 3). In consequence, these VPNs may be subject to tonic inhibition (from glu-lOLP) under constant light conditions and to phasic inhibition (from the Phasic VPN) upon an increase in Rh6-PRs dependent light intensity. Therefore, phasic inhibition from the Phasic VPN may potentially refine the temporal resolution of VPNs responses to increment of light intensity (ON response). The Phasic VPN is also under the control of glu-lOLP (Figure 3C), suggesting that it is in turn also subject to tonic inhibition.

An interesting aspect resulting from tonic inhibition of VPNs during constant light stimulation is what happens when the light intensity decreases (OFF response). Some neurons subject to tonic inhibition can emit more spikes when inhibition is lifted (Marder and Bucher, 2001; Hedwig 2016). If, via this mechanism, VPNs were to increase their firing rate upon a decrease in light intensity, they would be encoding an OFF response. Potentially all VPNs under tonic inhibition from glu-lOLP could have this rebound of activity after a light intensity decrease. Interestingly, two cholinergic VPNs (the nc-LaN 1 and 2) synapse onto the Phasic VPN that could then become more strongly activated by the OFF response (Figure 3D, Figure 1-figure supplement 2). As the Phasic VPN is inhibiting several VPNs (the 5^th^-LaN, the two nc-LaNs and PVL09), this could therefore create a second period of phasic inhibition allowing to maintain these VPNs OFF responses brief.

In conclusion, the connectivity of the third-order glutamatergic Phasic VPN putatively inhibitory suggests that it may refine the temporal resolution of VPNs ON and OFF responses through phasic inhibition.

### The visual circuit supports previous behavioral observations

In addition, this complete wiring diagram of the larval visual system can explain why in previous studies only Rh5-PRs appeared to be required for light avoidance while Rh6-PRs seemed dispensable in most experimental conditions (Keene et al., 2011; Kane et al., 2013; Humberg and Sprecher, 2017). When Rh6-PRs are mutated or functionally silenced, Rh5-PRs may still provide light absolute intensity information and the cholinergic inputs from nc-LaNs onto the glutamatergic Phasic VPN could inhibit the VPNs, computing on its own some response to changes in light intensity. Blocking the Phasic VPN activity in a Rh6-PRs depletion background would allow to study whether absolute light information is sufficient for visual navigation. Moreover, when Rh5-PRs are disabled, VPNs do not receive absolute light information while potentially being under tonic inhibition from the glu-lOLP, which therefore could shut down their activity. If glu-lOLP is indeed a provider of tonic inhibition, this would also suggest that, in most experimental conditions, ON and OFF responses alone are not sufficient to allow larval light avoidance but that it may need both a baseline activity of VPNs, provided by absolute light intensity information, and the modulations of this baseline activity in response to light intensity variations.

Only the Pdf-LaNs involved in circadian rhythm receive direct inputs from both PRs subtypes and can therefore still receive absolute light information in either PRs’ depletion condition. This broad light information integration capacity is in agreement with evidences that both PRs subtypes are sufficient to entrain the larval clock (Keene et al., 2011). While further reconstructions of their postsynaptic partners are required, the four Pdf-LaNs appear identical in sensory inputs, local connections and anatomy, raising the question of what such redundancy would allow.

### Comparisons between the larval eye and adult compound eye

With our data, similarities between the larval eye and the more complex compound eye of the adult fly, as well as with the vertebrate eye, are more and more striking (Figure 6). First these three visual systems have each two main types of photo-sensory neurons: Rh5-and Rh6-PRs for the *Drosophila* larva, inner and outer PRs for adult flies, cones and rods for vertebrates (Sprecher et al., 2007; Sprecher and Desplan, 2008; Friedrich, 2008; Sanes and Zipursky, 2010). Here we also confirmed that the LON was organized in layers (LONd, LONi, LONp) similarly as the adult flies optic lobe (lamina, medulla) and the vertebrate retina (outer and inner plexiform layers) (Sanes and Zipursky, 2010, Figure 6). Moreover, the mode of development of *Drosophila* larval and adult eyes reinforces similarities between Rh5-PRs with an inner-PR type and Rh6-PRs with an outer-PR type: in the adult R8-PR precursors are formed first to recruit outer PRs, likewise in the larva Rh5-PR precursors develop first and then recruit the Rh6-PRs, in both cases via the EGFR pathway. Also, development of both inner PRs and Rh5-PRs depend on the transcription factors *senseless* and *spalt* (Sprecher et al., 2007; Sprecher and Desplan, 2008; Mishra et al., 2013). Finally, the inputs from serotoninergic and octopaminergic/tyraminergic neurons onto the LON are also a shared feature with the adult visual system where serotonin has been linked to circadian rhythmicity (Yuan et al., 2005) and where visual cues during flight are modulated by octopaminergic inputs (Suver et al., 2012; Wasserman et al., 2015).

**Figure 6:**
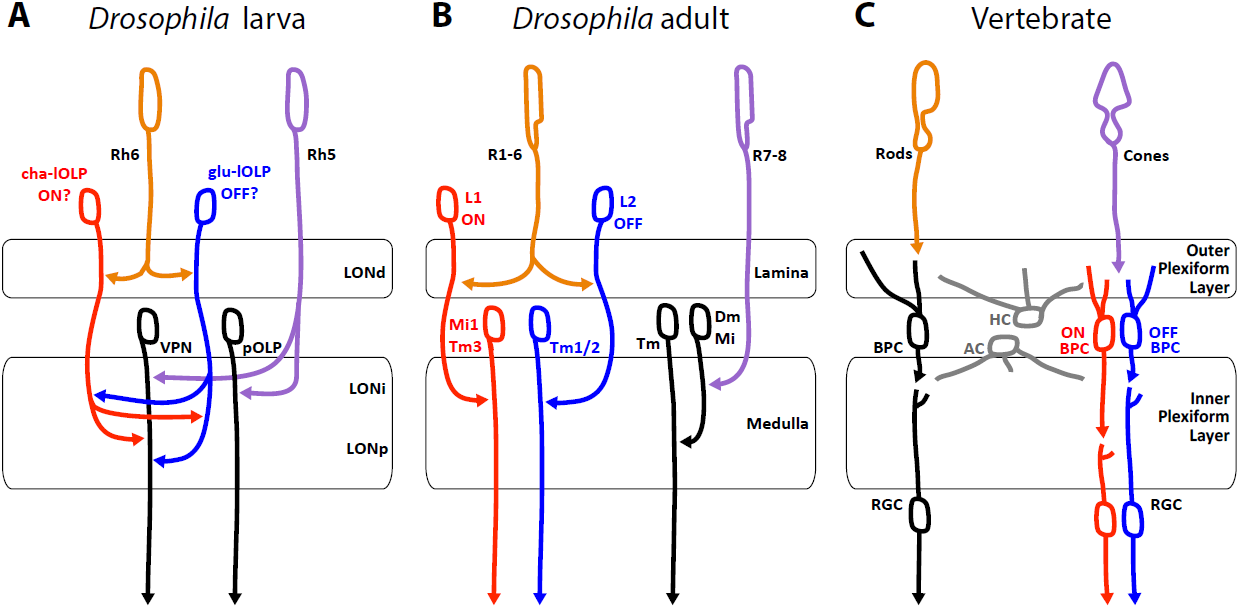
Comparison of the Drosophila larval visual circuit with the Drosophila adult compound eye and the vertebrate visual circuit. **A:** Larval visual circuit as described in this paper. Two main groups of VPNs receive input from Rh5-PRs (purple): one corresponds to the pOLP that only receives inputs from Rh5-PRs, whereas the second group (VPN) also receives inputs from cha and glu-lOLPs (red and blue) that are targets of Rh6-PRs (orange). We propose that cha and glu-lOLPs control light intensity increment and decrement (ON/OFF) detection respectively and transfer these information to the VPN group. **B:** Model of a single unit of the fly compound eye where R1-6 PRs (orange) are well known to be involved in contrast and motion detection whereas R7-8 PRs (purple) are involved in color sensing (Sanes and Zipursky, 2010; Clark and Demb, 2016, for reviews). In the lamina, R1-6 PRs make connections to the glutamatergic L1 neuron controlling the ON pathway (red) and to the cholinergic L2 neuron controlling the OFF pathway (blue). In the deeper medulla, L1 and L2 reach their targets (Mi1, Tm1/2), whereas R7-8 PRs connect to medullar neurons (Dm/Mi). **C:** Model of the vertebrate visual circuit (Sanes and Zipursky, 2010; Clark and Demb, 2016, for reviews). Cones (purple), which are also the colour sensors of the retina, connect to bipolar retinal cells (BPC) which constitute the ON or OFF pathways depending on the glutamate receptor they express (ON BPC and pathway in red, OFF BPC and pathway in blue). Rods (orange) also connect to BPC and control vision in dim light conditions. LONd, LONi and LONp: dorsal, intermediate and proximal larval optic neuropil. Mi: medulla intrinsic neurons; Tm: transmedulla neurons; Dm: dorsal medulla neurons. RGC: retina ganglion cells; HC: horizontal cells; AC: amacrine cells.

A functional comparison between the larval and the adult visual system emerge from the similar neurotransmitters expressed and the functions proposed for the larval VLNs cha-and glu-lOLP and the adult glutamatergic L1 and cholinergic L2 interneurons of the lamina (Figure 6A, B). In the adult fly, the outer R1-6 PRs connect to L1 and L2 interneurons that convey distinct responses to light increment (ON) and decrement (OFF) (Takemura et al., 2011). The adult PRs are histaminergic and induce hyperpolarization (inhibition) in both L1 and L2 (Dubs et al., 1981; Stuart, 1999; reviewed in Borst and Helmstaedter, 2015). Therefore upon light increment, PRs inhibit the glutamatergic L1, which results in the disinhibition of the downstream targets of L1 (ON response). In turn, upon light decrement PRs inhibit less the cholinergic L2, which results in the activation of the downstream targets of L2 (OFF response). However, the larval PRs are cholinergic (Yasuyama et al., 1995; Keene et al., 2011) and, at least for the Pdf-LaNs (Yuan et al., 2011), excite their targets activity in response of light. Therefore in larvae, the ON response would result from an increase of excitation from cha-lOLP onto VPNs when light intensity increases (via the increase of Rh6-PRs inputs) and could be kept transient by inhibition from the glu-lOLP (indirect presynaptic inhibition of Rh6-PRs inputs) and by phasic inhibition from the Phasic VPN. In turn, the OFF response would result from the disinhibition of VPNs from glu-lOLP inhibition when light intensity decreases and may also be kept transient by phasic inhibition from the Phasic VPN. Therefore, whereas the glutamatergic L1 conveys the ON response and the cholinergic L2 conveys the OFF response in adult flies, we propose that the glu-lOLP conveys the OFF response and the cha-lOLP conveys the ON response in larvae (Figure 6A, B). While ON/OFF responses in adult flies and vertebrates (Figure 6B,C) are involved in motion detection, such ability within one eye is achieved through downstream direction-sensitive cells that integrate information from several points in space (Clark and Demb, 2016). As larval eyes lack ommatidia such capacity seems unlikely, however the ON/OFF detection could already suffice for larval visual navigation (Kane et al., 2013; Klein et al., 2015; Schulze et al., 2015; Humberg and Sprecher, 2017).

### Comparison between the visual and the olfactory first-order processing centers

The LON neural network presents many similarities with the larval olfactory wiring diagram (Berck et al., 2016), favoring a potential common organizational origin of these sensory neuropils as suggested before (Strausfeld, 1989; Strausfeld et al., 2007). Odorant cues are perceived by olfactory receptor neurons that project to the antennal lobe where they contact olfactory projection interneurons and olfactory local interneurons. Sensory inputs in the antennal lobe are also segregated, not in layers like in visual circuits, but in olfactory receptor specific glomeruli (Fishilevich et al., 2005; Masuda-Nakagawa et al., 2009). Most olfactory receptor neurons and olfactory projection interneurons are cholinergic like PRs and VPNs (Keene et al., 2011; Yasuyama and Salvaterra 1999; Python and Stocker, 2002), and we find glutamatergic, potentially inhibitory, local interneurons in both systems (Berck et al., 2016). Reciprocal synapses between cha-lOLP and glu-lOLP in the visual circuit may be functionally equivalent to the connections between some olfactory receptor neurons and the glutamatergic Picky olfactory local interneuron 0 of the antennal lobe, suggesting that this reciprocal motif in the LON could indeed contribute to detecting changes in light intensity by computing the first derivative of the stimulus (Berck et al., 2016). Additionally, we can observe presynaptic and postsynaptic inhibition in both sensory systems while the strategies of implementation are somewhat different. Interestingly, we observed that most projection interneurons of both sensory systems may bring a mixture of absolute stimulus intensity and acceleration in stimulus intensity to the lateral horn and to the mushroom body calyx (Figure 3; Kim et al. 2015). Finally, both sensory systems are modulated by aminergic neurons inputs on specifics local interneurons. Interestingly, both the lOLPs VLNs and the olfactory Broad local interneurons Trio form feedback synapses onto the axonal termini of their respective bilateral octopaminergic/tyraminerigc neurons while this is not the case for serotonergic neurons (Berck et al., 2016).

### Concluding remarks

Identification of synaptic connectivity and neurotransmitter identity within the larval visual circuit allow us to formulate clear predictions on the response profile and function of individual network components. Based on the circuit map we suggest that the Rh6-PRs-VLNs pathway might be required for the detection of light intensity changes, whereas the Rh5-PRs-VPNs pathway could provide direct absolute light intensity information. In the future behavioral studies or physiological activity recording of visual circuit neurons will allow to add additional layers onto this functional map.

## Material and Methods

### ssTEM based neuronal reconstruction

In order to reconstruct the larval visual system, we used the serial-section transmission electron microscopy (ssTEM) volume of the entire nervous system of a first instar larva as described in Ohyama et al. (2015). Briefly, a 6-h-old *[iso] CantonS G1 x w1118* brain was 50nm serial cut and imaged at high TEM resolution. After images processing and compression, the whole dataset was stored on servers accessible by the web page CATMAID (Collaborative annotation Toolkit for Massive Amounts of Image Data, http://openconnecto.me/catmaid/, Saalfeld et al., 2009). The reconstruction was performed manually following the method used in Ohyama et al. (2015) and described in detail in Schneider-Mizell et al. (2016). All photoreceptor neurons were traced from the Bolwig nerve’s entrance in the ssTEM stack up to their terminals within the larval optic neuropil. The loss of eight 50-nm serial sections between frames 1103 and 1112 have made difficult to get all the neurons complete especially in the right hemisphere. Therefore, attempts to cross the gap for a neuron were validated with its contralateral homolog. We found 60 neurons in total, measuring 12.5 millimeters of cable and presenting 2090 presynaptic sites and 4414 postsynaptic sites. 7 tiny fragments (that amount to 0.018 millimeters of cable and 20 postsynaptic sites in total) could not be attached to full neuronal arbors. The reconstruction required 134 hours plus an additional 43 hours for proofreading.

### Fly strains

Flies were reared on standard cornmeal and molasses food in a 12h: 12h light-dark cycle at 25°C. We used the following strains for each subset of neurons (number of neurons of interest; neuron’s name): from the GMR Rubin GAL4 (R) and JRC split-Gal4 (SS0) collections: SS01740 (1; serotonergic SP2-1 neuron), SS02149 (2; octopaminergic/tyraminergic sVUM2 neurons), R72A10-GAL4 (3; OLPs), SS01724 (1; glutamatergic lOLP), SS01745 (1; projection OLP), R20C08-GAL4 and SS00671 (1; PVL09), R19C05-GAL4 (3; nc-LaNs and 5^th^-LaN; plus the 4 Pdf-LaNs are weakly expressed) and SS01777 (1; Phasic VPN). From Bloomington BDSC: OK371-GAL4 (VGluT-Gal4, #26160), and pJFRC29-10xUAS-IVS-myr::GFP-p10 (attP40) (referred as UAS-myr::GFP, #32198). Kind gift from B. Egger: w;; UAS-His2B-mRFP/TM3 (Mayer et al., 2005).

### Immunohistochemistry

All confocal stacks are from early third instar larvae. All identification of a neuron neurotransmitter expression was performed on first and third instars. Larvae were dissected 4 days after egg laying. Brain dissections where performed in cold phosphate-buffered saline (PBS, BioFroxx, 1X in dH2O). Brains were fixed in 3.7% formaldehyde in 1X PBS, 5mM MgCl_2_ (Merck), 0.5mM EGTA (Fluka) at room temperature for 25 minutes, except when using anti-DVGluT-antibody for which brains were fixed using Bouin’s solution for 5 minutes (picric acid/formaldehyde/glacial acetic acid in proportion 15/5/1, Daniels et al., 2008). Brains were stained according to previously described protocols (Sprecher et al., 2011) and mounted in DAPI-containing Vectashield (Vector laboratories). The following primary antibodies were used: rabbit anti-GFP (Sigma) and sheep anti-GFP (Serotec) (both 1:1000), mouse anti-ChAT (DHSB, 1:20 for neuropil marker and 1:5 for cell neurotransmitter identification), rabbit anti-serotonin (1:1000, Sigma, S5545), rabbit anti-DVGluT (1:5000, kind gift from A. DiAntonio (Daniels et al., 2008)) and rabbit anti-PER (1:1000, kind gift from R. Stanewsky (Gentile et al., 2013). For double ChAT / DVGluT staining brains were fixed following DVGluT protocol and mouse anti-ChAT was used at 1:2. The following secondary antibodies were used: donkey anti-sheep IgG Alexa Fluor 488, goat anti-rabbit IgG Alexa 488 and 647, goat anti-mouse IgG Alexa 488, 647 and 568 (all 1:200, Molecular Probes). Images were recorded using a Leica SP5 confocal microscope with a 63X 1.4 NA glycerol immersion objective and LAS software. Z-projections were made in Fiji (Software, NIH) and brightness adjustment with Adobe Photoshop®.

## Acknowledgements

We would like to sincerely thank Tim-Henning Humberg and our colleagues of the Sprecher lab, Akira Fushiki and Quan Yuan for fruitful discussions throughout the project. We thank the Developmental Studies Hybridoma Bank (DHSB), the Bloomington Drosophila Stock Center (BDSC), Aaron DiAntonio, Ralf Stanewsky and Boris Egger for providing fly lines and antibodies; Gerry Rubin for sharing GAL4 lines prior to publication. This work was supported by the Swiss National Science Foundation (CRSII3_136307 and 31003A_169993) and the European Research Council (ERC-2012-StG 309832-PhotoNaviNet) to S.G.S.

## Additional files

Figure 1 supplement 1

Figure 1 supplement 2

Figure 1 supplement 3

Figure 3 supplement animations

---

**Figure 1 - figure supplement 2:**
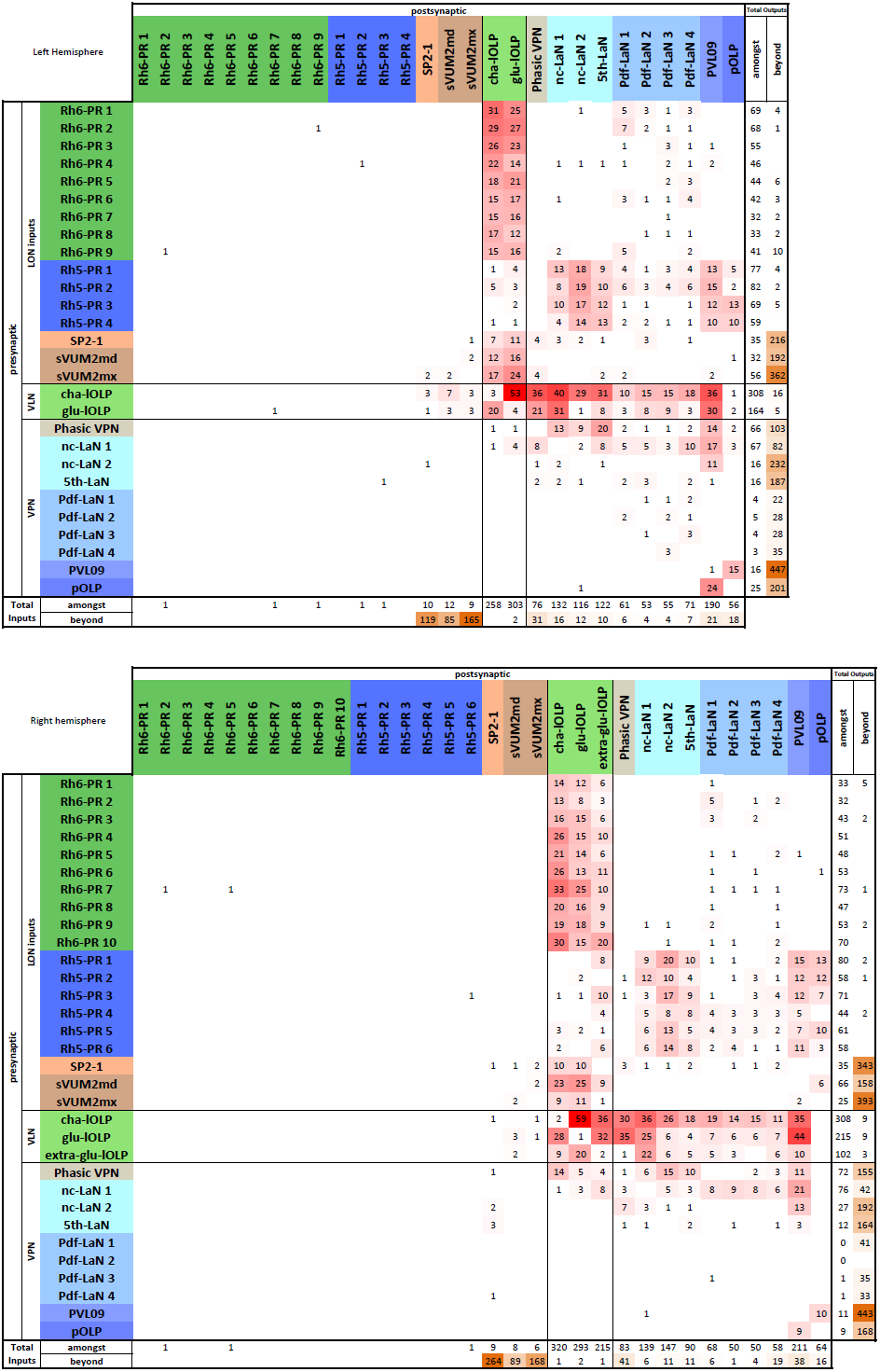
Complete synaptic connection matrices from both left (top) and right (lower) LON (data also provided in supplement files).

**Figure 1 - figure supplement 3:**
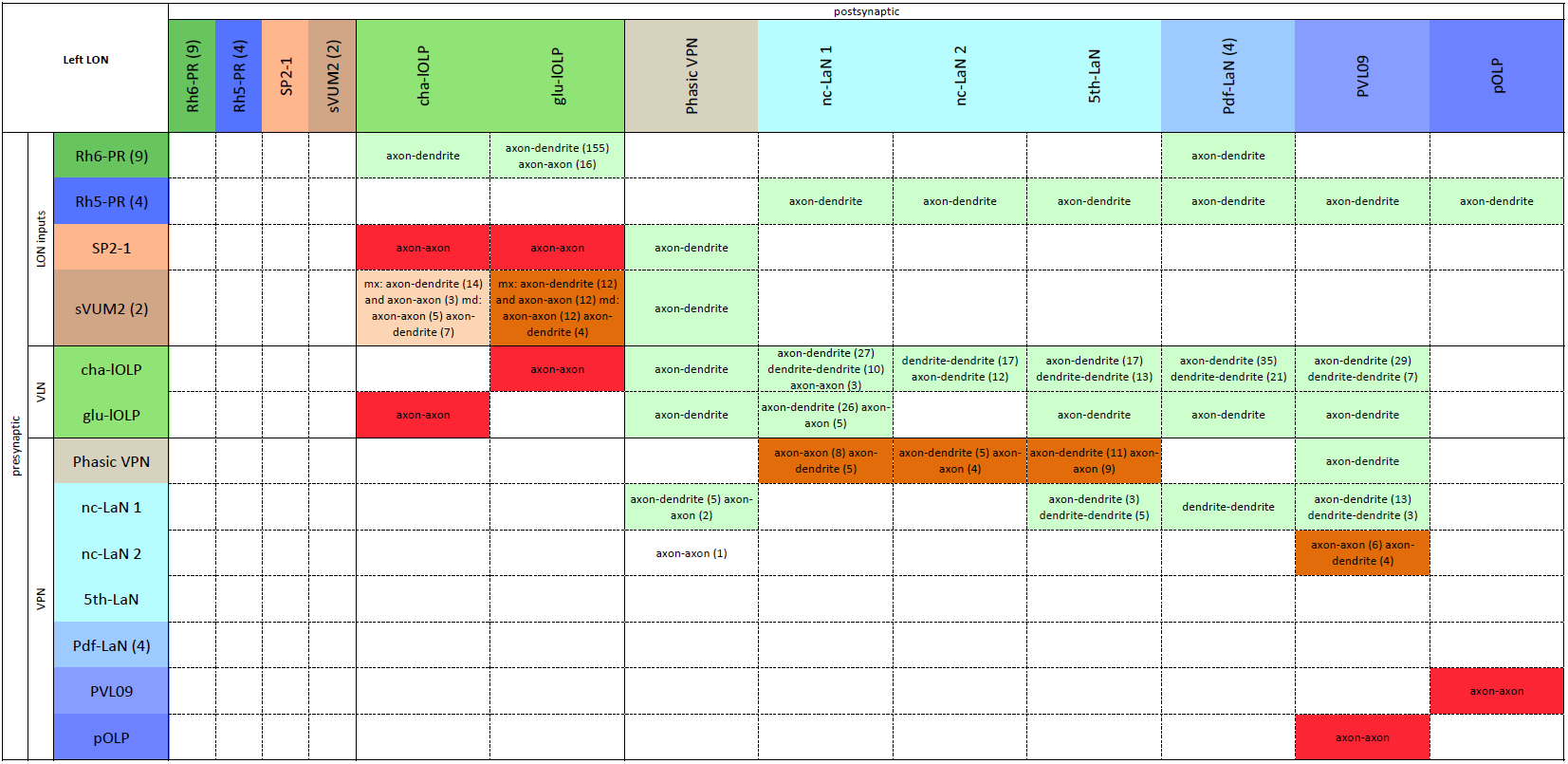
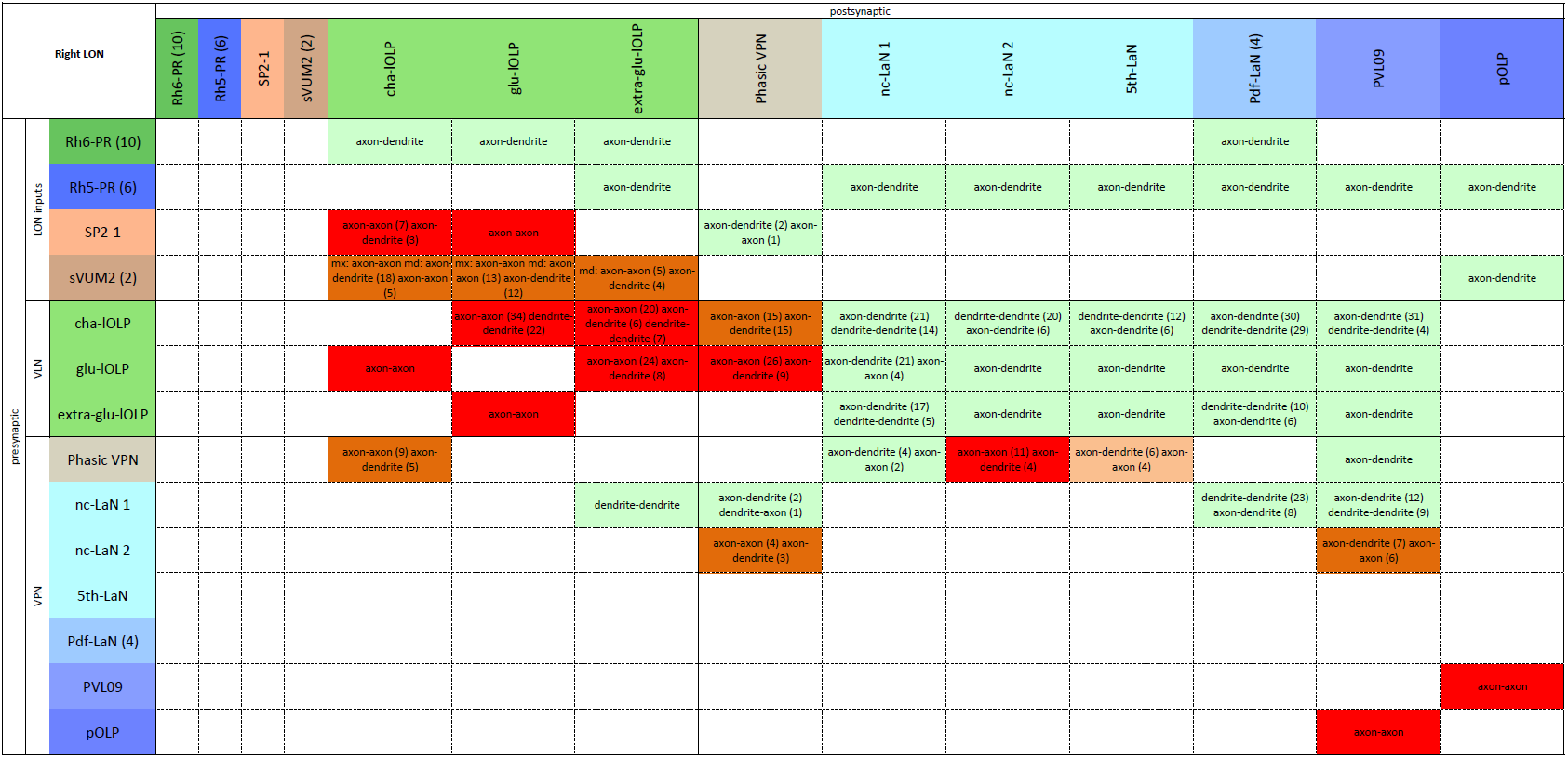
Main connection types for the right LON. In brackets: number of synapses when there is more than one type of connection (data also provided in supplement files).

**Figure 1 - figure supplement 4:**
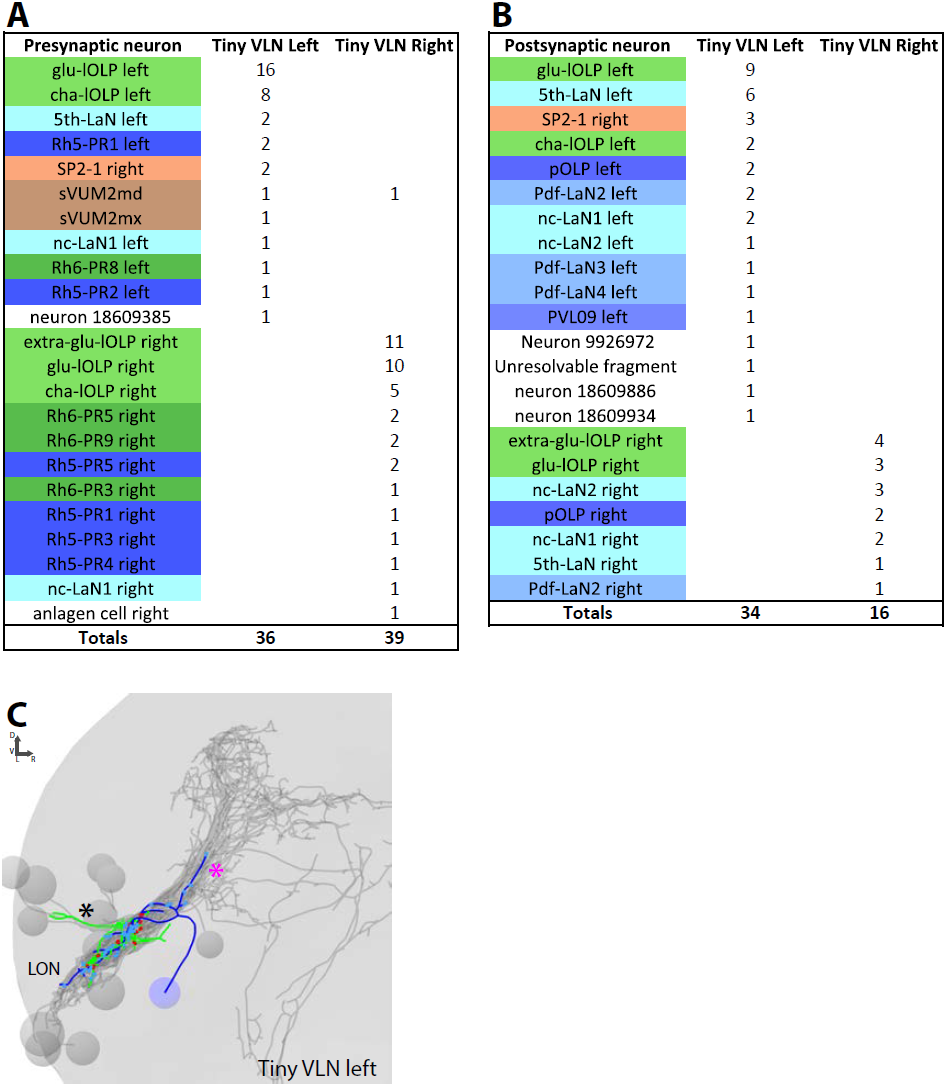
Connections and anatomy of the small third-order neuron Tiny VLN. **A:** The main inputs of both Tiny VLNs come from the lOLPs in particular the glu-lOLP (and extra-glu-lOLP in the right hemisphere). **B:** Tiny VLNs have few outputs, especially in the right hemisphere, but it seems that their main targets are back to the glu-lOLPs. **C:** 3D reconstruction of the left Tiny VLN with a medial-situated cell body, sparse connections in the LON and neurits entering different primary tracts (BLAd tract: black star; central optic tract: magenta star). Posterior view, dendrites in blue, axon in green, presynaptic sites in red, postsynaptic sites in cyan, other LON neurons in gray.

**Figure 1 - figure supplement 5:**
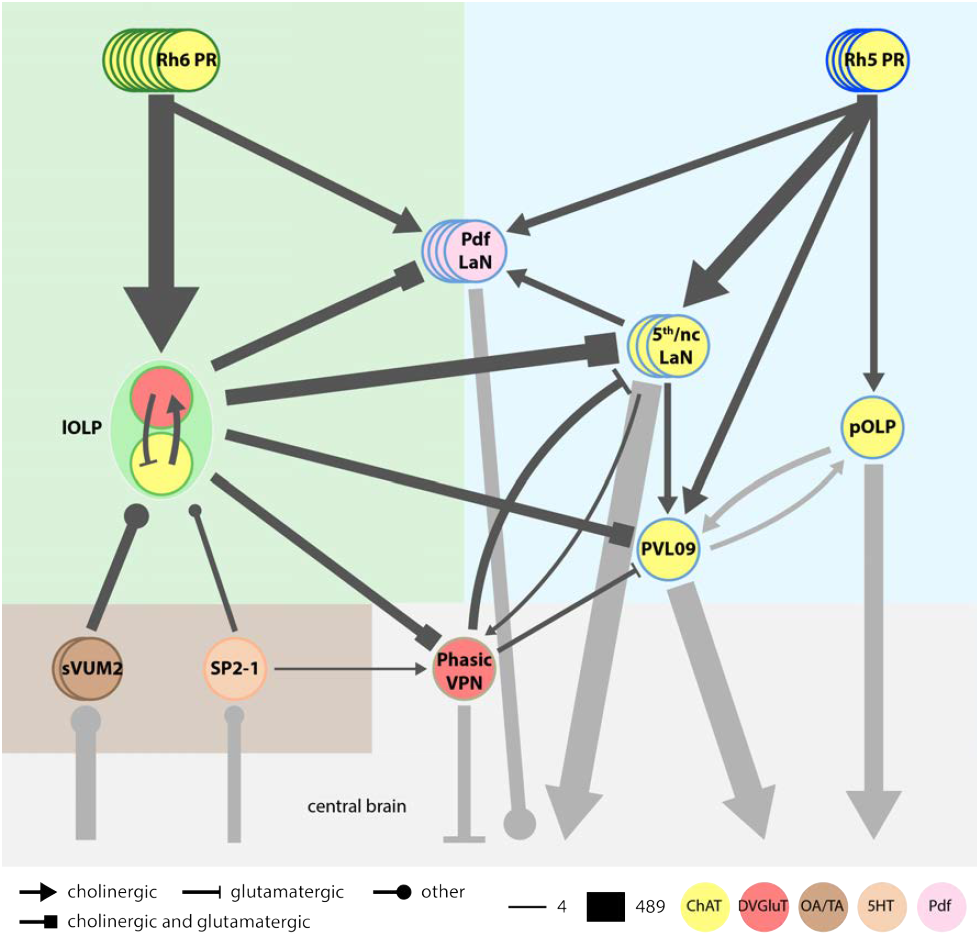
Model of the complete larval visual neural network. Each PRs subtypes have specific targets. Rh6-PRs (dark green) mainly contact the two lOLPs while Rh5-PRs (dark blue) contact VPNs. Only the Pdf-LaNs receive inputs from both PRs and therefore could be placed at the limit between both Rh6-PRs and Rh5-PRs pathways (green and blue areas). Cell circle outline color defines the neuron identity. Cell filled color represents the neurotransmitter/neuropeptide expression: yellow for cholinergic cells, red for glutamatergic cells, brown for octopaminergic/tyraminergic cells, light orange for serotonergic cell, pink for Pdf neuropeptide. All PRs neurons are cholinergic (Yasuyama et al., 1995; Keene et al., 2011). sVUM2 neurons (brown) are octopaminergic/tyraminergic. SP2-1 (orange) is serotonergic. One lOLP (light green) is cholinergic while the other one is glutamatergic. VPNs (shades of blue): pOLP and PVL09 are cholinergic; Pdf-LaNs express the Pdf neuropeptide and may co-express glycine (Frenkel et al., 2017); both nc-LaNs and the 5th-LaN are cholinergic. The third-order neuron Phasic VPN (light brown) is glutamatergic and, as it does not receive direct input from PRs, is not included in neither Rh6-PRs nor Rh5-PRs pathways nor with the aminergic modulatory neurons (brown area). For simplicity, the 5th-LaN and the two nc-LaNs are grouped together. Both lOLPs form a reciprocally connected pair that modulates almost all other VPNs. The Phasic VPN also modulates the 5th/nc-LaNs group as well as PVL09 while nc-LaN1 in particular positively reinforces Pdf-LaNs activity and both nc-LaNs connect the Phasic VPN. pOLP is the only VPN that is not modulated by other visual interneurons except for its strong reciprocal connections with PVL09 at their axons level. Black arrows represent connections within LON neurons while gray arrows represent connections beyond. Additional external inputs onto some VPNs are not represented here. Arrow thickness weighted by the square root of the number of synapses, arrow thickness scale shows minimum and maximum.

**Figure 2 - figure supplement 1:**
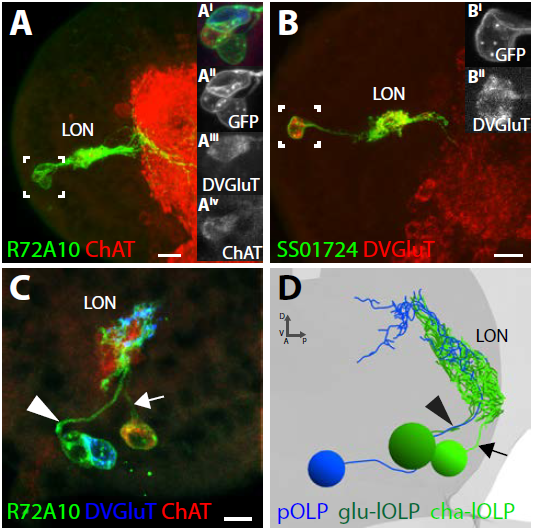
**A-C:** Confocal z-projections, dorsal view. A: R72A10>UAS-myr::GFP (green) showing the three OLPs (neuropil marker: ChAT, red) and close-up on the cells bodies (box) showing that one OLP is glutamatergic (**A^iii^**) and at least another one is clearly cholinergic (**A^iv^**). B: SS01724>UAS-myr::GFP (green) showing a lOLP with dense arborization within the LON, and reduced projections, which is glutamatergic (DVGluT in red) (**B^i^**, **B^ii^**close-up of the cell body (box)). **C:** R72A10>UAS-myr::GFP (green) close up on the three OLPs cell bodies where we can observe the strongly cholinergic cell (ChAT, red) sending its axon towards the LON via a different path (arrow) than the glutamatergic cell (DVGluT, blue) and the third cell (arrowhead). **D:** 3D reconstruction of the three OLPs in the left hemisphere of the ssTEM dataset where we could observe two cells sending their axons together to the LON (arrowhead), whereas the third one takes a separate path (arrow). Comparing **C** and **D** and based on their anatomy, the three OLPs can be distinguished in the EM (**D**): the projection OLP (pOLP, blue) and the glutamatergic lOLP (glu-lOLP, green) axons fasciculate together but not with the cholinergic lOLP (cha-lOLP, light green). Scale bars A, B: 10μm; C: 5μm.

**Figure 3 - figure supplement 1:**
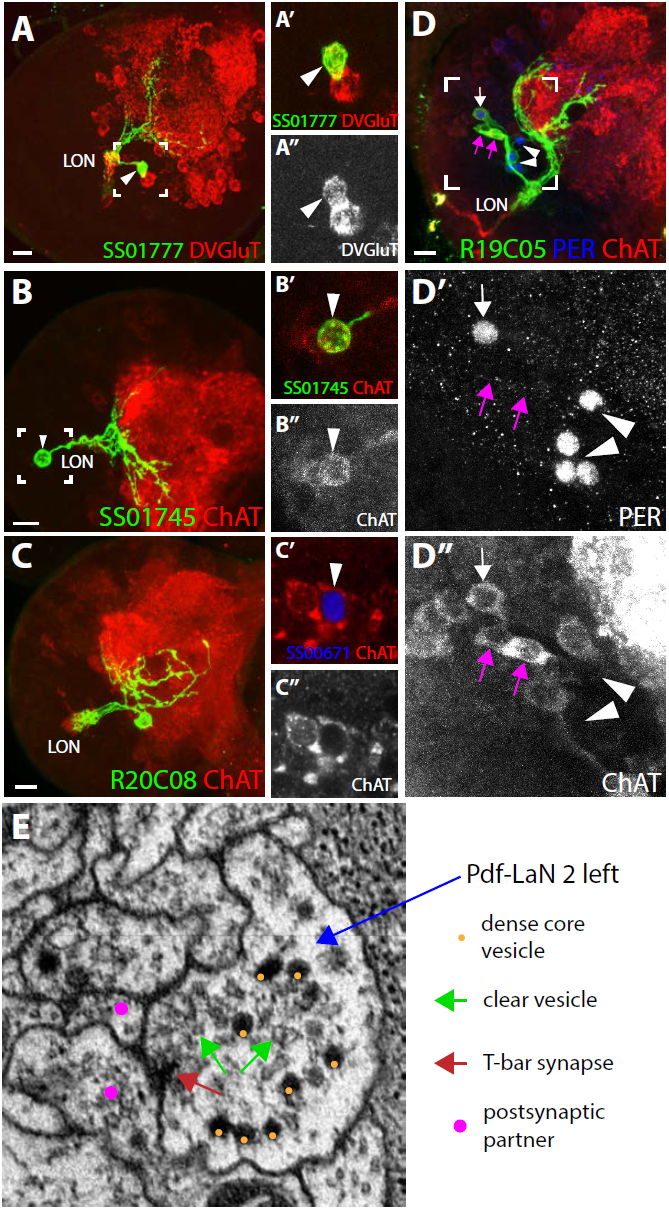
**A:** SS01777>UAS-myr::GFP (green) showing a stochastic single clone of the Phasic VPN that appears glutamatergic (arrowhead, DVGluT in red/white, **A′, A″** close up). **B:** SS01745>UAS-myr::GFP (green) showing the pOLP anatomy with weak arborization within the LON and deep projection into the neuropil. **B′, B″:** Close up on pOLP cell body showing a weak but clear cholinergic cell (arrow-head, ChAT in red/white). **C:** R20C08>UAS-myr::GFP (green) showing a single looping neuron with a strong overlap in the LON and having its cell body situated postero-ventro-laterally, corresponding to PVL09 (neuropil marker ChAT in red). **C′, C″:** Close up on a PVL09 cell body (SS0671>UAS-H2B-RFP, RFP in blue) that is cholinergic (ChAT, red, arrowhead). Single section. **D:** R19C05>UAS-myr::GFP (green) showing three cells among which only one is PER-positive (5th-LaN, white arrow) and two are PER-negatives (nc-LaNs, magenta arrow) (**D′**, PER in blue/white). All three cells are cholinergic (**D″**, ChAT in red/white). Four additional cells weakly covered by the Ga4 line and expressing PER correspond to the Pdf-LaNs (**D′**, arrowheads). Confocal z-projections, dorsal views, scale bars: A, B, C, D: 10μm. **E:** Electron microscopy view of a bouton rich in dense-core vesicles and clear vesicles in the Pdf-LaN 2 of the left hemisphere.

**Figure 3 - figure supplement 2:**
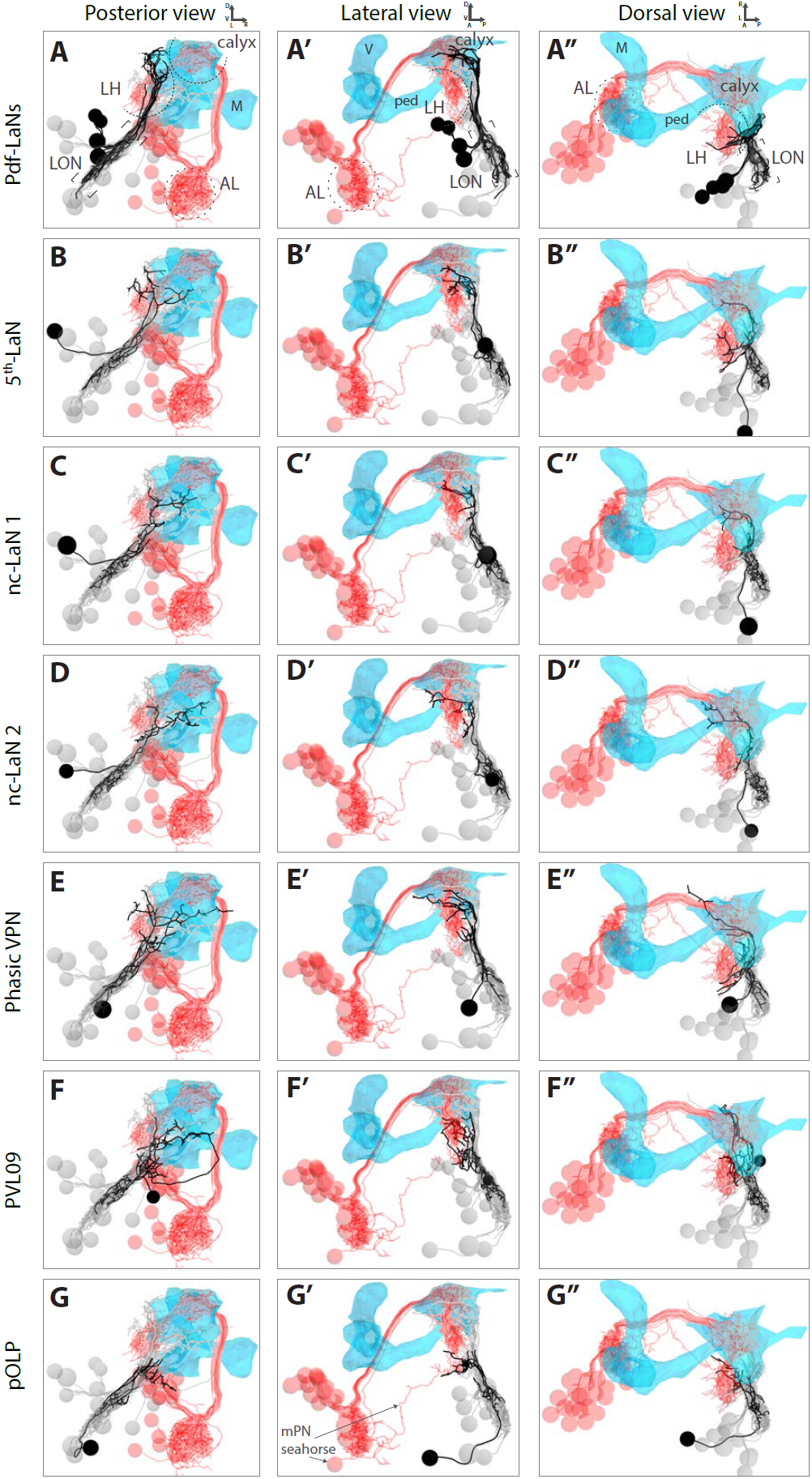
3D reconstructions of all VPNs from different views (posterior, lateral, dorsal) relative to the lateral horn (LH) shown by displaying olfactory projection interneurons (in red, AL: antennal lobe), and to the mushroom body (MB, blue mesh, V: vertical lobe, M: medial lobe, ped: peduncle). VPN of interest in black, other LON neurons in gray. **A-A″:** The Pdf-LaNs project above the LH. The 5th-LaN (**B-B″**), the nc-LaN 1 (**C-C″**), the nc-LaN 2 (**D-D″**) and the third-order neuron Phasic VPN (**E-E″**) project to the LH and MB calyx. **F-F″**: PVL09 projects to the LH and, after a long looping branch below the MB, project to the same region as pOLP. **G-G″:** pOLP projects like the multiglomerular olfactory projection neurons (mPN) Seahorse in the lower LH (two Pdf-LaNs were remove for these panels to unmask the lower LH). 3D animations of the rotating brain are provided in supplemental files.

**Figure 3 - figure supplement 3:**
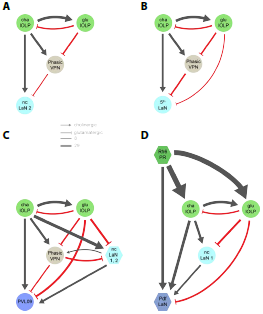
The nc-LaN 2 (**A**) and the 5th-LaN (**B**) are controlled in similar fashion by the two lOLPs and the Phasic VPN. **C:** PVL09 activity is shaped by the same motives from both the lOLPs and the Phasic VPN and is additionally controlled by a second level of coherent feedforward loops (FFL) from the two nc-LaNs. **D:** Unlike other VPNs, the Pdf-LaNs receive direct inputs from Rh6-PRs. The Pdf-LaNs are downstream of two interlocked coherent FFLs from the Rh6-PRs via cha-lOLP and nc-LaN 1. Left hemisphere, hexagons represent group of cells, circles represent single cell, arrow thickness weighted by the square root of the number of synapses, arrow thickness scale shows minimum and median.

**Figure 4 - figure supplement 1:**
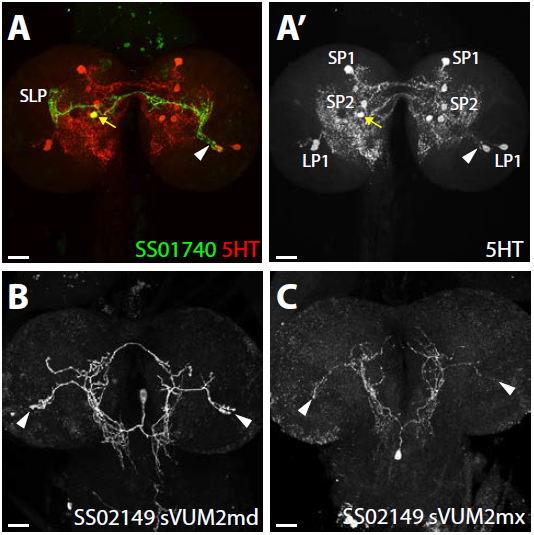
**A:** Confocal z-projection of SS01740>UAS-myr::GFP (green) showing a stochastic single clone of SP2-1 neuron (yellow arrow) innervating the contralateral LON (arrowhead, serotonin (5-HT) in red). **A′:** anti-5-HT channel shows the three 5-HT clusters from the lower and superior protocerebrum (LP and SP) and innervation in the LON (arrowhead). **B-C:** Confocal z-projection of SS02149>UAS-myr::GFP (white) showing stochastic single clone expression of sVUM2md (**B**) and sVUM2mx (**C**) innervating both LON (arrowheads). Scale bars: 20μm.

**Figure 5 - figure supplement 1:**
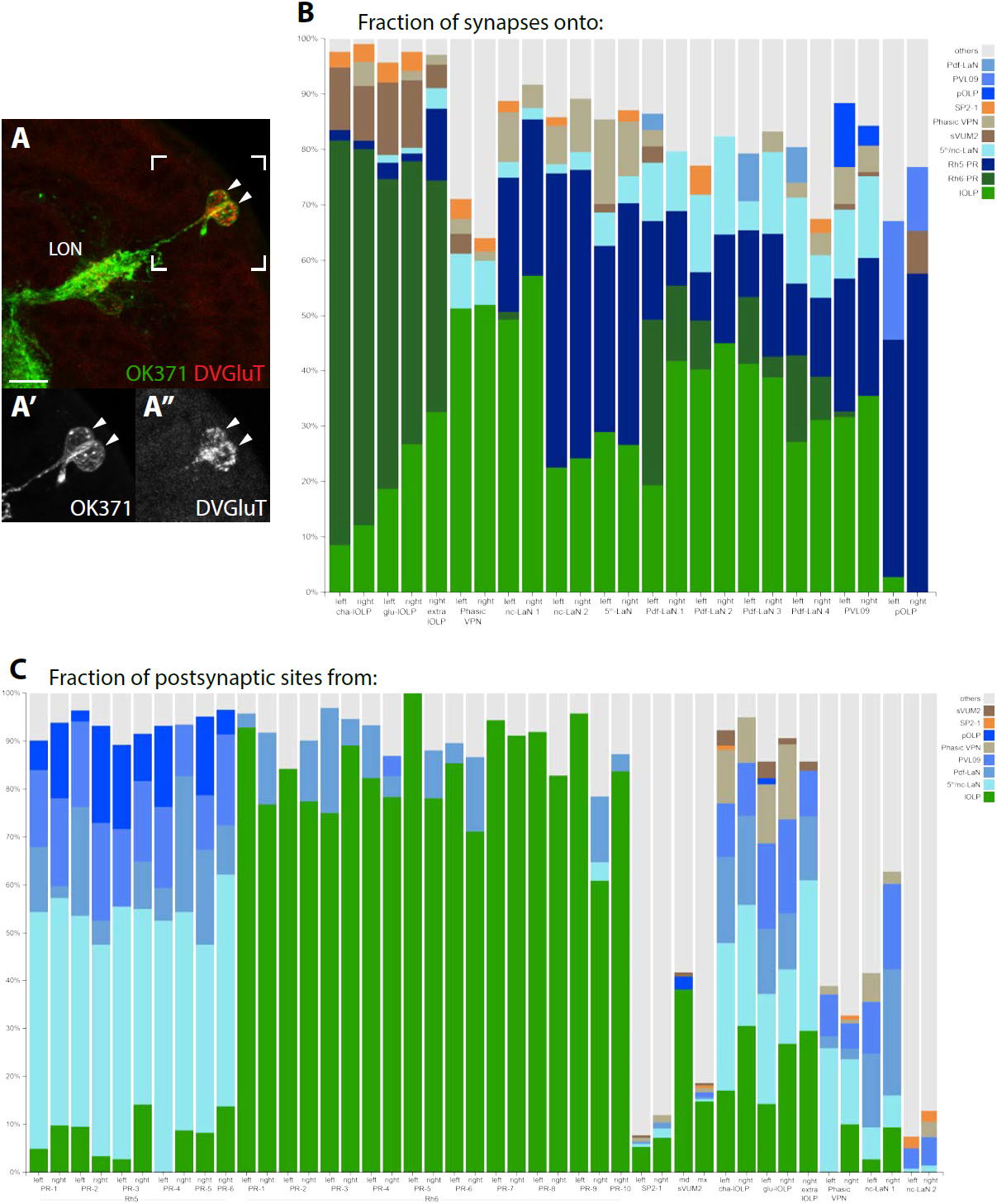
**A:** Confocal z-projection of OK371-Gal4>UAS-myr::GFP (green) stained with anti-DVGluT (red) with insets onto two cells bodies in the OLP region and projecting into the LON and which are both DVGluT positive (arrow heads, close up **A′**, **A″**). Dorsal view, scale bar: 10 μm. **B:** Percentage of synapses onto all visual interneurons of both hemispheres (two bars per neuron except for the extra-lOLP of the right hemisphere). lOLPs main inputs are from the Rh6-PRs. The Phasic VPN does not receives from either PRs but main inputs come from the lOLPs. All other VPNs receive from Rh5-PRs and only the Pdf-LaNs receives from both PRs-subtypes. Except for pOLP, the lOLPs connections represent an important fraction of each neuron inputs. The Phasic VPN inputs are most significant on the two nc-LaNs and the 5th-LaN. **C:** Percentage of synapses from all PRs, aminergic modulatory neurons and visual interneurons with significant roles in local processing (lOLPs, Phasic VPN and nc-LaNs). Additional PRs from the right hemisphere have clear connectivity profiles of either Rh5 or Rh6 subtypes. Strong involvement of the aminergic modulatory neurons in other neural circuits is visible.

## References

Alon, U. (2007). Network motifs: theory and experimental approaches. Nat Rev Genet, 8(6), 450–461. doi:10.1038/nrg2102

Baines, R. A., & Bate, M. (1998). Electrophysiological development of central neurons in the Drosophila embryo. J Neurosci, 18(12), 4673–4683.

Berck, M. E., Khandelwal, A., Claus, L., Hernandez-Nunez, L., Si, G., Tabone, C. J., … Cardona, A. (2016). The wiring diagram of a glomerular olfactory system. Elife, 5. doi:10.7554/eLife.14859

Borst, A., & Helmstaedter, M. (2015). Common circuit design in fly and mammalian motion vision. Nat Neurosci, 18(8), 1067–1076. doi:10.1038/nn.4050

Burrows, M. (1996). The Neurobiology of an Insect Brain: Oxford University Press.

Clark, D. A., & Demb, J. B. (2016). Parallel Computations in Insect and Mammalian Visual Motion Processing. Curr Biol, 26(20), R1062–R1072. doi:10.1016/j.cub.2016.08.003

Dacks, A. M., Green, D. S., Root, C. M., Nighorn, A. J., & Wang, J. W. (2009). Serotonin modulates olfactory processing in the antennal lobe of Drosophila. J Neurogenet, 23(4), 366–377. doi:10.3109/01677060903085722

Daniels, R. W., Gelfand, M. V., Collins, C. A., & DiAntonio, A. (2008). Visualizing glutamatergic cell bodies and synapses in Drosophila larval and adult CNS. J Comp Neurol, 508(1), 131–152. doi:10.1002/cne.21670

Delgado, R., Barla, R., Latorre, R., & Labarca, P. (1989). L-glutamate activates excitatory and inhibitory channels in Drosophila larval muscle. FEBS Lett, 243(2), 337–342.

Dubs, A., Laughlin, S. B., & Srinivasan, M. V. (1981). Single photon signals in fly photoreceptors and first order interneurones at behavioral threshold. J Physiol, 317, 317–334.

Eichler, K., Li, F., Litwin-Kumar, A., Park, Y., Andrade, I., Schneider-Mizell, C. M., Saumweber, T., Huser, A., Bonnery, D., Gerber, B. et al. (under revision). The complete wiring diagram of a high-order learning and memory center, the insect mushroom body.

Fishilevich, E., Domingos, A. I., Asahina, K., Naef, F., Vosshall, L. B., & Louis, M. (2005). Chemotaxis behavior mediated by single larval olfactory neurons in Drosophila. Curr Biol, 15(23), 2086–2096. doi:10.1016/j.cub.2005.11.016

Frenkel, L., Muraro, N. I., Beltran Gonzalez, A. N., Marcora, M. S., Bernabo, G., Hermann-Luibl, C., … Ceriani, M. F. (2017). Organization of Circadian Behavior Relies on Glycinergic Transmission. Cell Rep, 19(1), 72–85. doi:10.1016/j.celrep.2017.03.034

Friedrich, M. (2008). Opsins and cell fate in the Drosophila Bolwig organ: tricky lessons in homology inference. Bioessays, 30(10), 980–993. doi:10.1002/bies.20803

Gepner, R., Mihovilovic Skanata, M., Bernat, N. M., Kaplow, M., & Gershow, M. (2015). Computations underlying Drosophila photo-taxis, odor-taxis, and multi-sensory integration. Elife, 4. doi:10.7554/eLife.06229

Gerber, B., Scherer, S., Neuser, K., Michels, B., Hendel, T., Stocker, R. F., & Heisenberg, M. (2004). Visual learning in individually assayed Drosophila larvae. J Exp Biol, 207(Pt 1), 179–188.

Gong, Z. (2009). Behavioral dissection of Drosophila larval phototaxis. Biochem. Biophys. Res. Commun. 382, 395–399. doi: 10.1016/j.bbrc.2009.03.033

Hamasaka, Y., Rieger, D., Parmentier, M. L., Grau, Y., Helfrich-Forster, C., & Nassel, D. R. (2007). Glutamate and its metabotropic receptor in Drosophila clock neuron circuits. J Comp Neurol, 505(1), 32–45. doi:10.1002/cne.21471

Hassan, J., Iyengar, B., Scantlebury, N., Rodriguez Moncalvo, V., & Campos, A. R. (2005). Photic input pathways that mediate the Drosophila larval response to light and circadian rhythmicity are developmentally related but functionally distinct. J Comp Neurol, 481(3), 266–275. doi:10.1002/cne.20383

Hedwig, B. G. (2016). Sequential Filtering Processes Shape Feature Detection in Crickets: A Framework for Song Pattern Recognition. Front Physiol, 7, 46. doi:10.3389/fphys.2016.00046

Helfrich-Forster, C. (1997). Development of pigment-dispersing hormone-immunoreactive neurons in the nervous system of Drosophila melanogaster. J Comp Neurol, 380(3), 335–354.

Humberg, T.-H., & Sprecher, S. G. (2017). Age-and Wavelength-Dependency of Drosophila Larval Phototaxis and Behavioral Responses to Natural Lighting Conditions. Frontiers in Behavioral Neuroscience, 11(66). doi:10.3389/fnbeh.2017.00066

Huser, A., Rohwedder, A., Apostolopoulou, A. A., Widmann, A., Pfitzenmaier, J. E., Maiolo, E. M., … Thum, A. S. (2012). The serotonergic central nervous system of the Drosophila larva: anatomy and behavioral function. PLoS One, 7(10), e47518. doi:10.1371/journal.pone.0047518

Jovanic, T., Schneider-Mizell, C. M., Shao, M., Masson, J. B., Denisov, G., Fetter, R. D., … Zlatic, M. (2016). Competitive Disinhibition Mediates Behavioral Choice and Sequences in Drosophila. Cell, 167(3), 858–870 e819. doi:10.1016/j.cell.2016.09.009

Kane, E. A., Gershow, M., Afonso, B., Larderet, I., Klein, M., Carter, A. R., … Samuel, A. D. (2013). Sensorimotor structure of Drosophila larva phototaxis. Proc Natl Acad Sci U S A, 110(40), E3868–3877. doi:10.1073/pnas.1215295110

Kaneko, M., Helfrich-Forster, C., & Hall, J. C. (1997). Spatial and temporal expression of the period and timeless genes in the developing nervous system of Drosophila: newly identified pacemaker candidates and novel features of clock gene product cycling. J Neurosci, 17(17), 6745–6760.

Keene, A. C., Mazzoni, E. O., Zhen, J., Younger, M. A., Yamaguchi, S., Blau, J., … Sprecher, S. G. (2011). Distinct visual pathways mediate Drosophila larval light avoidance and circadian clock entrainment. J Neurosci, 31(17), 6527–6534. doi:10.1523/JNEUROSCI.6165-10.2011

Kim, A. J., Lazar, A. A., & Slutskiy, Y. B. (2015). Projection neurons in Drosophila antennal lobes signal the acceleration of odor concentrations. Elife, 4. doi:10.7554/eLife.06651

Klein, M., Afonso, B., Vonner, A. J., Hernandez-Nunez, L., Berck, M., Tabone, C. J., … Samuel, A. D. (2015). Sensory determinants of behavioral dynamics in Drosophila thermotaxis. Proc Natl Acad Sci U S A, 112(2), E220–229. doi:10.1073/pnas.1416212112

Linster, C., & Smith, B. H. (1997). A computational model of the response of honey bee antennal lobe circuitry to odor mixtures: overshadowing, blocking and unblocking can arise from lateral inhibition. Behav Brain Res, 87(1), 1–14.

Liu, W. W., & Wilson, R. I. (2013). Glutamate is an inhibitory neurotransmitter in the Drosophila olfactory system. Proc Natl Acad Sci U S A, 110(25), 10294–10299. doi:10.1073/pnas.1220560110

Mahr, A., & Aberle, H. (2006). The expression pattern of the Drosophila vesicular glutamate transporter: a marker protein for motoneurons and glutamatergic centers in the brain. Gene Expr Patterns, 6(3), 299–309. doi:10.1016/j.modgep.2005.07.006

Majeed, Z. R., Abdeljaber, E., Soveland, R., Cornwell, K., Bankemper, A., Koch, F., & Cooper, R. L. (2016). Modulatory Action by the Serotonergic System: Behavior and Neurophysiology in Drosophila melanogaster. Neural Plast, 2016, 7291438. doi:10.1155/2016/7291438

Malpel, S., Klarsfeld, A., & Rouyer, F. (2002). Larval optic nerve and adult extra-retinal photoreceptors sequentially associate with clock neurons during Drosophila brain development. Development, 129(6), 1443–1453.

Marder, E., & Bucher, D. (2001). Central pattern generators and the control of rhythmic movements. Curr Biol, 11(23), R986–996.

Masuda-Nakagawa, L. M., Gendre, N., O’Kane, C. J., & Stocker, R. F. (2009). Localized olfactory representation in mushroom bodies of Drosophila larvae. Proc Natl Acad Sci U S A, 106(25), 10314–10319. doi:10.1073/pnas.0900178106

Mayer, B., Emery, G., Berdnik, D., Wirtz-Peitz, F., & Knoblich, J. A. (2005). Quantitative analysis of protein dynamics during asymmetric cell division. Curr Biol, 15(20), 1847–1854. doi:10.1016/j.cub.2005.08.067

Mazzoni, E. O., Desplan, C., & Blau, J. (2005). Circadian pacemaker neurons transmit and modulate visual information to control a rapid behavioral response. Neuron, 45(2), 293–300. doi:10.1016/j.neuron.2004.12.038

Mishra, A. K., Tsachaki, M., Rister, J., Ng, J., Celik, A., & Sprecher, S. G. (2013). Binary cell fate decisions and fate transformation in the Drosophila larval eye. PLoS Genet, 9(12), e1004027. doi:10.1371/journal.pgen.1004027

Nagel, K. I., Hong, E. J., & Wilson, R. I. (2015). Synaptic and circuit mechanisms promoting broadband transmission of olfactory stimulus dynamics. Nat Neurosci, 18(1), 56–65. doi:10.1038/nn.3895

Ohyama, T., Schneider-Mizell, C. M., Fetter, R. D., Aleman, J. V., Franconville, R., Rivera-Alba, M., … Zlatic, M. (2015). A multilevel multimodal circuit enhances action selection in Drosophila. Nature, 520(7549), 633–639. doi:10.1038/nature14297

Olsen, S. R., Bhandawat, V., & Wilson, R. I. (2010). Divisive normalization in olfactory population codes. Neuron, 66(2), 287–299. doi:10.1016/j.neuron.2010.04.009

Olsen, S. R., & Wilson, R. I. (2008). Lateral presynaptic inhibition mediates gain control in an olfactory circuit. Nature, 452(7190), 956–960. doi:10.1038/nature06864

Python, F., & Stocker, R. F. (2002). Immunoreactivity against choline acetyltransferase, gammaaminobutyric acid, histamine, octopamine, and serotonin in the larval chemosensory system of Dosophila melanogaster. J Comp Neurol, 453(2), 157–167. doi:10.1002/cne.10383

Randel, N., Asadulina, A., Bezares-Calderon, L. A., Veraszto, C., Williams, E. A., Conzelmann, M., … Jekely, G. (2014). Neuronal connectome of a sensory-motor circuit for visual navigation. Elife, 3. doi:10.7554/eLife.02730

Randel, N., Shahidi, R., Veraszto, C., Bezares-Calderon, L. A., Schmidt, S., & Jekely, G. (2015). Inter-individual stereotypy of the Platynereis larval visual connectome. Elife, 4, e08069. doi:10.7554/eLife.08069

Ready, D. F., Hanson, T. E., & Benzer, S. (1976). Development of the Drosophila retina, a neurocrystalline lattice. Dev Biol, 53(2), 217–240.

Rodriguez Moncalvo, V. G., & Campos, A. R. (2005). Genetic dissection of trophic interactions in the larval optic neuropil of Drosophila melanogaster. Dev Biol, 286(2), 549–558. doi:10.1016/j.ydbio.2005.08.030

Rodriguez Moncalvo, V. G., & Campos, A. R. (2009). Role of serotonergic neurons in the Drosophila larval response to light. BMC Neurosci, 10, 66. doi:10.1186/1471-2202-10-66

Roy, B., Singh, A. P., Shetty, C., Chaudhary, V., North, A., Landgraf, M., … Rodrigues, V. (2007). Metamorphosis of an identified serotonergic neuron in the Drosophila olfactory system. Neural Dev, 2, 20. doi:10.1186/1749-8104-2-20

Rybak, J., Talarico, G., Ruiz, S., Arnold, C., Cantera, R., & Hansson, B. S. (2016). Synaptic circuitry of identified neurons in the antennal lobe of Drosophila melanogaster. J Comp Neurol, 524(9), 1920–1956. doi:10.1002/cne.23966

Saalfeld, S., Cardona, A., Hartenstein, V., & Tomancak, P. (2009). CATMAID: collaborative annotation toolkit for massive amounts of image data. Bioinformatics, 25(15), 1984–1986. doi:10.1093/bioinformatics/btp266

Saalfeld, S., Fetter, R., Cardona, A., & Tomancak, P. (2012). Elastic volume reconstruction from series of ultra-thin microscopy sections. Nat Methods, 9(7), 717–720. doi:10.1038/nmeth.2072

Sanes, J. R., & Zipursky, S. L. (2010). Design principles of insect and vertebrate visual systems. Neuron, 66(1), 15–36. doi:10.1016/j.neuron.2010.01.018

Sawin-McCormack, E. P., Sokolowski, M. B., & Campos, A. R. (1995). Characterization and genetic analysis of Drosophila melanogaster photobehavior during larval development. J Neurogenet, 10(2), 119–135.

Schlegel, P., Texada, M. J., Miroschnikow, A., Schoofs, A., Huckesfeld, S., Peters, M., … Pankratz, M. J. (2016). Synaptic transmission parallels neuromodulation in a central food-intake circuit. Elife, 5. doi:10.7554/eLife.16799

Schneider-Mizell, C. M., Gerhard, S., Longair, M., Kazimiers, T., Li, F., Zwart, M. F., … Cardona, A. (2016). Quantitative neuroanatomy for connectomics in Drosophila. Elife, 5. doi:10.7554/eLife.12059

Schulze, A., Gomez-Marin, A., Rajendran, V. G., Lott, G., Musy, M., Ahammad, P., … Louis, M. (2015). Dynamical feature extraction at the sensory periphery guides chemotaxis. Elife, 4. doi:10.7554/eLife.06694

Selcho, M., Pauls, D., El Jundi, B., Stocker, R. F., & Thum, A. S. (2012). The role of octopamine and tyramine in Drosophila larval locomotion. J Comp Neurol, 520(16), 3764–3785. doi:10.1002/cne.23152

Selcho, M., Pauls, D., Han, K. A., Stocker, R. F., & Thum, A. S. (2009). The role of dopamine in Drosophila larval classical olfactory conditioning. PLoS One, 4(6), e5897. doi:10.1371/journal.pone.0005897

Selcho, M., Pauls, D., Huser, A., Stocker, R. F., & Thum, A. S. (2014). Characterization of the octopaminergic and tyraminergic neurons in the central brain of Drosophila larvae. J Comp Neurol, 522(15), 3485–3500. doi:10.1002/cne.23616

Sprecher, S. G., Cardona, A., & Hartenstein, V. (2011). The Drosophila larval visual system: high-resolution analysis of a simple visual neuropil. Dev Biol, 358(1), 33–43. doi:10.1016/j.ydbio.2011.07.006

Sprecher, S. G., & Desplan, C. (2008). Switch of rhodopsin expression in terminally differentiated Drosophila sensory neurons. Nature, 454(7203), 533–537. doi:10.1038/nature07062

Sprecher, S. G., Reichert, H., & Hartenstein, V. (2007). Gene expression patterns in primary neuronal clusters of the Drosophila embryonic brain. Gene Expr Patterns, 7(5), 584–595. doi:10.1016/j.modgep.2007.01.004

Strausfeld, N. J. (1989). Insect Vision and Olfaction: Common Design Principles of Neuronal Organization. In R. N. Singh & N. J. Strausfeld (Eds.), Neurobiology of Sensory Systems (pp. 319–353). Boston, MA: Springer US.

Strausfeld, N. J., Sinakevitch, I., & Okamura, J. Y. (2007). Organization of local interneurons in optic glomeruli of the dipterous visual system and comparisons with the antennal lobes. Dev Neurobiol, 67(10), 1267–1288. doi:10.1002/dneu.20396

Stuart, A. E. (1999). From fruit flies to barnacles, histamine is the neurotransmitter of arthropod photoreceptors. Neuron, 22(3), 431–433.

Suver, M. P., Mamiya, A., & Dickinson, M. H. (2012). Octopamine neurons mediate flight-induced modulation of visual processing in Drosophila. Curr Biol, 22(24), 2294–2302. doi:10.1016/j.cub.2012.10.034

Takemura, S. Y., Karuppudurai, T., Ting, C. Y., Lu, Z., Lee, C. H., & Meinertzhagen, I. A. (2011). Cholinergic circuits integrate neighboring visual signals in a Drosophila motion detection pathway. Curr Biol, 21(24), 2077–2084. doi:10.1016/j.cub.2011.10.053

Takemura, S. Y., Xu, C. S., Lu, Z., Rivlin, P. K., Parag, T., Olbris, D. J., … Scheffer, L. K. (2015). Synaptic circuits and their variations within different columns in the visual system of Drosophila. Proc Natl Acad Sci U S A, 112(44), 13711–13716. doi:10.1073/pnas.1509820112

Tix, S., Minden, J. S., & Technau, G. M. (1989). Pre-existing neuronal pathways in the developing optic lobes of Drosophila. Development, 105(4), 739–746.

Tobin, W. F., Wilson, R. I., & Lee, W.-C. A. (2017). Wiring variations that enable and constrain neural computation in a sensory microcircuit. bioRxiv. doi:10.1101/097659

Varshney, L. R., Chen, B. L., Paniagua, E., Hall, D. H., & Chklovskii, D. B. (2011). Structural properties of the Caenorhabditis elegans neuronal network. PLoS Comput Biol, 7(2), e1001066. doi:10.1371/journal.pcbi.1001066

von Essen, A. M., Pauls, D., Thum, A. S., & Sprecher, S. G. (2011). Capacity of visual classical conditioning in Drosophila larvae. Behav Neurosci, 125(6), 921–929. doi:10.1037/a0025758

Vosshall, L. B., & Stocker, R. F. (2007). Molecular architecture of smell and taste in Drosophila. Annu Rev Neurosci, 30, 505–533. doi:10.1146/annurev.neuro.30.051606.094306

Wasserman, S. M., Aptekar, J. W., Lu, P., Nguyen, J., Wang, A. L., Keles, M. F., … Frye, M. A. (2015). Olfactory neuromodulation of motion vision circuitry in Drosophila. Curr Biol, 25(4), 467–472. doi:10.1016/j.cub.2014.12.012

Wilson, R. I., & Laurent, G. (2005). Role of GABAergic inhibition in shaping odor-evoked spatiotemporal patterns in the Drosophila antennal lobe. J Neurosci, 25(40), 9069–9079. doi:10.1523/JNEUROSCI.2070-05.2005

Wolf, H., & Burrows, M. (1995). Proprioceptive sensory neurons of a locust leg receive rhythmic presynpatic inhibition during walking. J Neurosci, 15(8), 5623–5636.

Wu, M., Nern, A., Williamson, W. R., Morimoto, M. M., Reiser, M. B., Card, G. M., & Rubin, G. M. (2016). Visual projection neurons in the Drosophila lobula link feature detection to distinct behavioral programs. Elife, 5. doi:10.7554/eLife.21022

Yamanaka, N., Romero, N. M., Martin, F. A., Rewitz, K. F., Sun, M., O’Connor, M. B., et al. (2013). Neuroendocrine control of Drosophila larval light preference. Science 341, 1113–1116. doi: 10.1126/science.1241210

Yasuyama, K., Kitamoto, T., & Salvaterra, P. M. (1995). Localization of choline acetyltransferase-expressing neurons in the larval visual system of Drosophila melanogaster. Cell Tissue Res, 282(2), 193–202.

Yasuyama, K., & Meinertzhagen, I. A. (2010). Synaptic connections of PDF-immunoreactive lateral neurons projecting to the dorsal protocerebrum of Drosophila melanogaster. J Comp Neurol, 518(3), 292–304. doi:10.1002/cne.22210

Yasuyama, K., & Salvaterra, P. M. (1999). Localization of choline acetyltransferase-expressing neurons in Drosophila nervous system. Microsc Res Tech, 45(2), 65–79. doi:10.1002/(SICI)1097-0029(19990415)45:2<65::AID-JEMT2>3.0.CO;2-0

Yuan, Q., Lin, F., Zheng, X., & Sehgal, A. (2005). Serotonin modulates circadian entrainment in Drosophila. Neuron, 47(1), 115–127. doi:10.1016/j.neuron.2005.05.027

Yuan, Q., Xiang, Y., Yan, Z., Han, C., Jan, L. Y., & Jan, Y. N. (2011). Light-induced structural and functional plasticity in Drosophila larval visual system. Science, 333(6048), 1458–1462. doi:10.1126/science.1207121

